# Evolutionarily Conserved Transcription Factors Drive the Oxidative Stress Response in *Drosophila*

**DOI:** 10.1101/735126

**Authors:** Sarah M. Ryan, Kaitie Wildman, Briseida Oceguera-Perez, Scott Barbee, Nathan T. Mortimer, Alysia D. Vrailas-Mortimer

**Author notes:** denotes equal contribution (Sarah M. Ryan, Kaitie Wildman). denotes equal contribution and co-corresponding authors (Nathan T. Mortimer, Alysia D. Vrailas-Mortimer). Corresponding authors: Alysia D. Vrailas-Mortimer, Illinois State University, School of Biological Sciences, Science Laboratory Building, Normal, IL 61790, Nathan T. Mortimer, Illinois State University, School of Biological Sciences, Science Laboratory Building, Normal, IL 61790.

## Abstract

As organisms are constantly exposed to the damaging effects of oxidative stress through both environmental exposure as well as internal metabolic processes, they have evolved a variety of mechanisms to cope with this stress. One such mechanism is the highly conserved p38 MAPK (p38K) pathway, which is known to be to post-translationally activated in response to oxidative stress resulting in the activation of downstream antioxidant targets. However, little is known about the role of p38K transcriptional regulation in response to oxidative stress. Therefore, we analyzed the p38K gene family across the genus *Drosophila* to identify conserved regulatory elements. We find that oxidative stress exposure results in increased p38K protein levels in multiple *Drosophila* species and is associated with increased oxidative stress resistance. We also find that the p38Kb genomic locus includes conserved binding sites for the AP-1 and lola-PT transcription factors. Accordingly, over-expression of these transcription factors in *D. melanogaster* is sufficient to induce transcription of p38Kb and enhances resistance to oxidative stress. We further find that the presence of a lola-PT binding site in the p38Kb locus of a given species is predictive of the species’ survival in response to oxidative stress. Through our comparative genomics approach, we have identified biologically relevant transcription factor binding sites that regulate the expression of p38Kb and are associated with resistance to oxidative stress. These findings reveal a novel mode of regulation for p38K genes and suggests that transcription may play as important a role in p38K mediated stress responses as post-translational modifications.

**Significance Statement:** Organisms encounter a variety of environmental stresses such as oxidative stress throughout their lifetime. Therefore, organisms have evolved a number of mechanisms to combat these stresses. In order to understand how these mechanisms evolved, we have compared the genomes of a diverse set of species across the genus *Drosophila* to examine the p38 MAPK stress response gene family. Our analysis was able to successfully predict transcription factors that not only regulate our target gene, p38Kb, but do so under different conditions to ensure an appropriate stress response. Therefore, we find that in addition to post-translational regulation, transcriptional regulation of signaling pathways may also play an important role in how organisms are able to adapt to stressful environments or respond to stress conditions as they arise. Furthermore, our comparative genomics approach may be utilized to identify transcriptional regulators of other highly conserved signaling pathways.

## Introduction

Organisms encounter oxidative stress, which damages a variety of cellular structures, both through external environmental exposure and from the internal release of oxygen radicals during ATP synthesis. Thus, organisms have evolved a variety of mechanisms to counteract it. One such mechanism is the highly conserved p38 MAP Kinase (p38K) pathway, which consists of a phosphorylation cascade of upstream kinases that leads to the dual phosphorylation of the p38K protein on a TXY motif. This activated p38K can phosphorylate target proteins in the cytoplasm and can also translocate to the nucleus where it can phosphorylate downstream transcription factors to induce the expression of antioxidant genes (*1–3*), allowing the cell to appropriately respond to the oxidative stress.

Though much is known about the role of post-translational regulation in the p38K mediated oxidative stress response, the transcriptional regulation of p38K during stress conditions is not well understood. In order to address this question, we have utilized a comparative genomics approach using the genus *Drosophila* to identify conserved transcription factor binding sites that play a role in mediating p38K gene expression during oxidative stress.

In the fruit fly *Drosophila melanogaster,* there are three p38K genes: p38Ka, p38Kb, and p38Kc. Loss of any one of these p38K genes is viable, however, p38Ka p38Kb double knockouts are lethal (*4–9*), suggesting that these genes are at least partially redundant. Both p38Ka and p38Kb play a role in resistance to heat shock, starvation, and oxidative stress (*4,9,10*). Furthermore, both of these genes play a role in the immune response with p38Ka regulating the gut DUOX system (*11*), while p38Kb is involved in bacterial phagocytosis (*5*) and antiviral immunity (*12*). In addition, these genes have unique roles in the body. p38Ka plays a role in cardiac function (*13*), while p38Kb is a regulator of lifespan, locomotor functions, and circadian rhythms (*4,14*). Interestingly, p38Kc has distinct functions in the cell in regulating innate immunity through the *Ddc* pathway (*15*) and intestinal oxidative stress and lipid homeostasis (*16*). The three p38K genes have been hypothesized to have arisen through gene duplication, and the differences in function suggest that evolutionary pressures may be playing a role in the divergence of these genes. As these genes play differing roles in the cell, they may have distinct genetic regulatory elements that control the expression of p38K during different developmental or homeostatic cellular responses.

In order to identify these potential regulatory elements, we utilized the sequenced genomes of 22 species with known evolutionary relationships within the *Drosophila* genus to identify conserved transcription factor binding sites upstream of the p38K family genes. We find that diverse species of *Drosophila* express high levels of p38K protein during oxidative stress, and that there are conserved AP-1 and lola-PT transcription factor binding sites that regulate p38Kb expression under normal and oxidative stress conditions. The levels of p38K expression and the presence of these transcription factor binding sites are also predictors of resistance to oxidative stress. These data suggest that transcriptional regulation of p38Kb may play as important a role in oxidative stress resistance and may boost the effectiveness of the more well-described post-translational regulation of p38Kb in regulating stress responses.

## Materials and Methods

### Sequences

Sequences corresponding to MAPK family genes were downloaded from FlyBase (flybase.org) and NCBI. Accession numbers are given in Table S1 for DNA sequences and Table S2 for protein sequences.

### Phylogenetic trees

Phylogenetic trees for the MAPK family were constructed using amino acid sequences. Sequences were aligned using the ClustalW algorithm with the Gonnet Protein Weight Matrix implemented in MEGA (version 5.2.2, (*17*)), and all positions containing gaps and missing data were eliminated. Initial trees were obtained using the Neighbor-Joining method. The phylogenetic relationships among the sequences were inferred using the maximum likelihood method based on the Jones-Taylor-Thornton model (*18*), and the trees with the highest log likelihood are shown. Each tree was tested with 1000 bootstrap replications (*19*), and the given values are the percentage of trees in which the associated sequences clustered together.

Phylogenetic trees for the *Drosophila* p38K gene family and ITS2 were constructed using nucleic acid sequences. Sequences were aligned using the ClustalW algorithm with the IUB DNA Weight Matrix implemented in MEGA, and all positions containing gaps and missing data were eliminated. Pairwise distances were estimated using the Maximum Composite Likelihood approach and initial trees were obtained using the Neighbor-Joining method. The phylogenetic relationships among the sequences were inferred using the maximum likelihood method based on the Tamura-Nei model (*20*), and the trees with the highest likelihood are shown. Each tree was tested with 1000 bootstrap replications (*19*), and the given values are the percentage of trees in which the associated sequences clustered together.

The output from these analyses were used to draw trees using FigTree v. 1.4.

### Chromosome synteny

The regions of interest from the Muller B and E elements across *Drosophila* species were accessed through GBrowse on FlyBase. Synteny was determined manually.

### Genetic conservation analysis

Pairwise dN and dS values for p38K family genes across the *Drosophila* species were computed in MEGA (v.5.2.2) using the Nei-Gojobori model. Values were plotted in R.

### Drosophila stocks

*Drosophila melanogaster* OregonR, y[1] w[*]; P{w[+mC]=UAS-lola.4.7}3, and P{w[+mC]=UAS-lola.4.7}1, w[*] were obtained from the Bloomington Drosophila Stock Center, Indiana University. w-;;MHC-GAL4 was used as reported in (*4*) and w-;UAS-Jun; UAS-Fos is described in (*21*). *Drosophila pseudoobscura* (strain 14011-0121.94), *Drosophila virilis* (strain 15010-1051.87), *Drosophila simulans* (strain 14021-0251.195), *Drosophila yakuba* (strain 14021-0261.00), *Drosophila mauritiana* (strain 14021-0241.151), and *Drosophila ananassae* (strain 14024-0371.13) were obtained from the Drosophila Species Stock Center, University of California San Diego.

### Predicting transcription factor binding sites

Transcription factor binding sites were predicted using iMotifs (*22*). The TomTom Motif Comparison Tool v4.9.1 (MEME web server, (*23*)) compared the resulting predicted sites to all known *Drosophila* transcription factor binding sites. The size of the letter corresponds to the program’s confidence at specific positions.

### qRTPCR

*D. melanogaster* lines were reared at 25°C in a 12hr:12hr light:dark cycle on standard fly food media. Virgin female flies were collected at eclosion and fed standard fly food media and aged for 1 week. Flies were then transferred to food mixed with 20mM paraquat (Sigma). Food was changed every 3-4 days. Surviving flies were collected when 30% of controls had died. Thoraxes from 6 flies were dissected and pooled. RNA was extracted using Trizol (Thermofisher Scientific) and reverse transcribed using SuperScript IV (Thermofisher Scientific). qRTPCR was performed using SYBR green (Thermofisher Scientific). Primers designed to amplify p38Kb or the housekeeping gene Arp88 were utilized. To compare p38Kb expression across genotypes, the ΔCt value for each replicate was calculated by subtracting the Ct value of Arp88 from the Ct value of p38Kb. We tested the hypothesis that AP-1 and Lola-PT overexpression would increase the amount of p38Kb transcript using one-tailed Student’s t and Wilcoxon signed rank tests (implemented in R).

### Lifespan

AP-1 and lola-PT flies and outcrossed controls were reared at 25°C in a 12hr:12hr light:dark cycle on standard fly food media. Flies were collected at eclosion and lifespan was assayed daily. Food was changed every 3-4 days. Differences in survival were assayed using the Log-Rank test. Pairwise comparisons between samples were made using the Pairwise Log-Rank test and p-values were adjusted using the false discovery rate. These tests were performed using the survival package in R.

### Paraquat Survival

We find that the efficacy of the paraquat was batch dependent (Table S3). Therefore, all experiments were performed with biological replicates and controls all on the same batch of paraquat. Virgin females from each species were collected at eclosion and transferred to fly food media mixed with 20mM paraquat (Sigma). *D. simulans* and *D. pseudoobscura* were reared at 21°C in a 12hr:12hr light:dark cycle. *D. melanogaster, D. virilis*, *D. yakuba*, *D. ananassae,* and *D. mauritiana* were reared at 25°C in a 12hr:12hr light:dark cycle on standard fly food media. Food was changed every 3-4 days. Survival was scored daily.

AP-1 and lola-PT flies and outcrossed controls were reared at 25°C in a 12hr:12hr light:dark cycle on standard fly food media. Virgin female flies were collected at eclosion and aged 1 week and then transferred to standard fly food media mixed with either 10mM or 20mM paraquat. Food was changed every 3-4 days. Survival was scored daily. Differences in survival were assayed using the Log-Rank test. Pairwise comparisons between samples were made using the Pairwise Log-Rank test and p-values were adjusted using the false discovery rate. These tests were performed using the survival package in R.

### Immunoblots

Three thoraxes from adult female flies where homogenized in 1X Laemmli buffer. Immunoblots were performed as described in (*4*). Membranes were probed with mouse anti-total p38 MAPK (1:1000, Upstate), rabbit phospho p38 MAPK (1:1000, Cell Signaling Technologies), and rabbit anti-alpha tubulin (1:5000) and either goat anti-mouse HRP 1:20,000 (Jackson Labs) or goat anti-rabbit HRP 1:40,000 (Jackson Labs). Membranes were then developed using SuperSignal West Femto kit (ThermoFisher) or Pierce ECL (ThermoFisher) and exposed on autoradiography film. All immunoblots were performed at least in triplicate. Densitometry was performed using Adobe Photoshop. For each replicate, the densitometry value of each p38K band was normalized to its corresponding alpha tubulin control. These normalized values were then fit to linear regression models with each blot treated as a separate blocking factor and compared using ANOVA (implemented in R).

### Interaction analysis

To test for a correlation between body size and survival on paraquat, we calculated the Pearson’s product-moment correlation between body size of female flies (approximated by thorax length, (*24*)) and the median survival following paraquat exposure. In an attempt to uncouple the phenotypic correlation from phylogenetic relationships, we used the phylogenetically independent contrasts approach (as implemented in the caper R package) to calculate the Spearman rho correlation between body size and median survival contrasts. To test for a correlation between p38K protein levels and survival on paraquat, we calculated the Spearman rho correlation between phylogenetically independent contrasts for the change in p38K protein levels in flies fed paraquat and the median survival following paraquat exposure. Analyses were performed in R and correlation data were plotted using ggplot2 in R.

To test the relationship between p38K protein levels, paraquat exposure and the presence of predicted transcription factor binding sites, we used a model in which paraquat exposure and site presence were independent variables and p38K level was the dependent variable. These data were compared using the Analysis of Variance of Aligned Rank Transformed Data method (implemented using the ARTool package in R). Interaction plots were generated using ggplot2 in R.

To test the relationship between the presence of predicted transcription factor binding sites and survival on paraquat, the data were compared using the Analysis of Variance of Aligned Rank Transformed Data method (ARTool package in R). Data were plotted using ggplot2 in R.

## Results

### Analysis of MAPK family genes across taxa

As p38K is closely related to the other the MAPK family members ERK and JNK, we first analyzed the relationship between the three MAPK families: ERK, JNK, and p38K across taxa (Tables S4-S6) as well as within a subset of the sequenced *Drosophila* species that are representative of the genus. While humans have multiple ERK and JNK family members, *D. melanogaster* only has one ortholog for each of these families (*rolled* and *basket*, respectively). In addition, vertebrates have four p38K genes while *D. melanogaster* have three p38K orthologs (*p38Ka*, *p38Kb*, and *p38Kc*). We find that all three MAPK families emerged very early in animal evolution before the split between vertebrates and invertebrates (Figure 1A). In addition, the ERK and JNK families evolved from a more recent ancestral gene, whereas the p38K family diverged earlier (Figure 1A). Within the p38K family, the three *D. melanogaste*r p38K genes are not orthologous to the distinct vertebrate p38K genes, suggesting that the family members have evolved since the split between vertebrates and invertebrates (Figure 1A). Analysis of a subset of *Drosophila* species reveals that these relationships are also conserved across the genus, with the species having single orthologs of ERK and JNK and multiple members of the p38K family (Figure 1B). This analysis further reveals that p38Kc is more recently diverged from p38Ka and p38Kb (Figure 1B). This is particularly interesting as in *D. melanogaste*r, the p38Kb gene resides on the second chromosome, while both p38Ka and p38Kc reside in close proximity on the third chromosome, suggesting that p38Ka and p38Kc may be the result of a tandem gene duplication.

**Figure 1.**
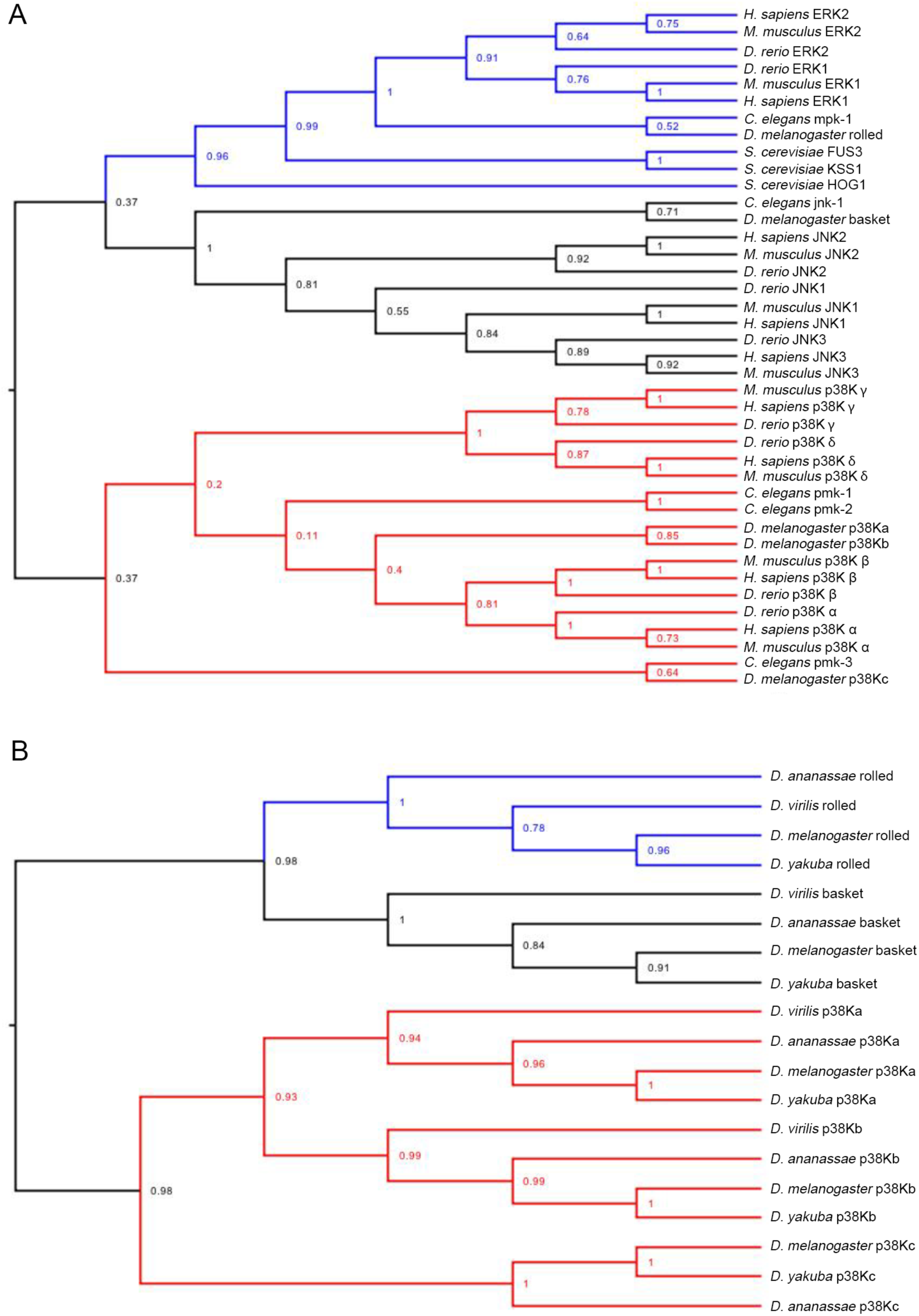
Phylogenetic Analysis of the MAPK Family. A) The evolutionary relationship between the ERK1/2 (blue), JNK (black), and p38 (red) MAPK genes from *Homo sapiens*, *Mus musculus*, *Danio rerio*, *Drosophila melanogaster*, *Caenorhabditis elegans,* and *Saccharomyces cerevisiae*. Note that HOG1 is the *Saccharomyces cerevisiae* homologue of both JNK and p38. B) The evolutionary relationship between the ERK1/2 (blue), JNK (black), and p38 (red) MAPK genes from representative species in the genus *Drosophila*.

### Comparison of p38K genes in Drosophila

In order to identify conserved genetic regulatory elements, we next compared the genomic locus of the p38K genes across 22 sequenced *Drosophila* species. We find that across species each p38K gene forms a distinct cluster (Figure S1). All three of the p38K genes mostly cluster along taxonomic lines as determined by molecular phylogeny (Figure 2A). Interestingly, unlike the other *Drosophila* species, *D. pseudoobscura* has two p38Kb genes (Figure 2C), suggesting a recent duplication of p38Kb in this species. As not all of the 22 sequenced *Drosophila* species have fully annotated genomes, we focused on 12 species that are fully sequenced and annotated to analyze the genomic architecture of the p38K loci. We find that the synteny of the p38Ka and p38Kc genes are highly conserved (Figure 3A), with their proximity suggesting that p38Kc most likely arose from a tandem duplication event of p38Ka, after the divergence of the willistoni group (Figures 2D and 3A). Furthermore, in *D. persimilis* p38Kc is truncated as compared to the p38Kc gene in other *Drosophila* species leading to a shortened p38Kc protein. Interestingly, *D. persimilis* also has three additional genes (GL24143, GL24139, GL24140) that have appeared near the p38Kc locus, disrupting its close proximity to p38Ka (Figure 3A). Both *D. sechellia* and *D. grimshawi* have independent duplications of a neighboring gene (CG6178), and *D. grimshawi* also has a new gene (GH19091) inserted into this region.

**Figure 2.**
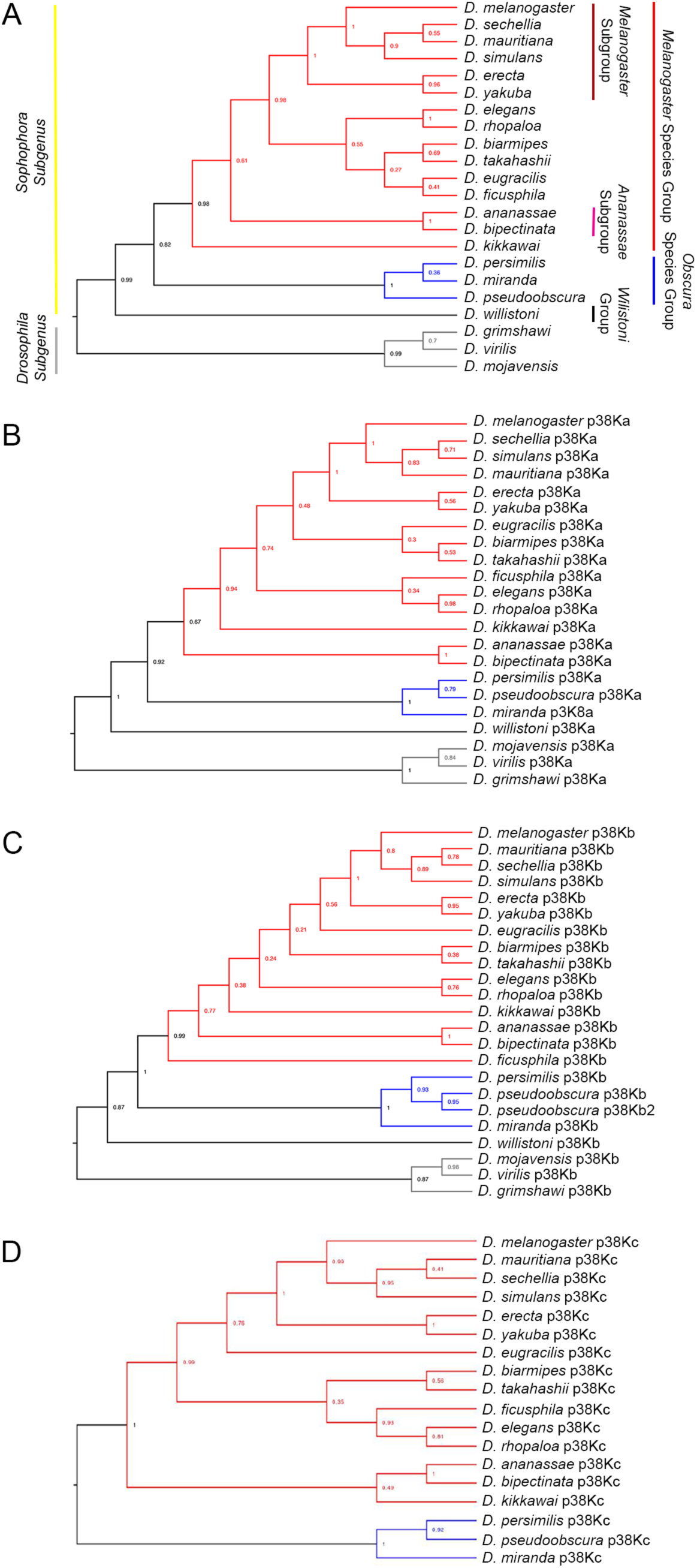
Phylogenetic Analysis of each p38K gene in *Drosophila*. The evolutionary relationships of the A) ITS2, B) p38Ka, C) p38Kb, and D) p38Kc genes in 22 sequenced species of *Drosophila*. Red denotes the melanogaster species group, blue denotes the obscura group, black denotes the willistoni group, and grey represents the *Drosophila* subgenus. Note that *D. pseudoobscura* is the only species with 2 copies of the p38Kb gene. p38Kc arose after the split of the obscura group from *D. willistoni*.

**Figure 3.**
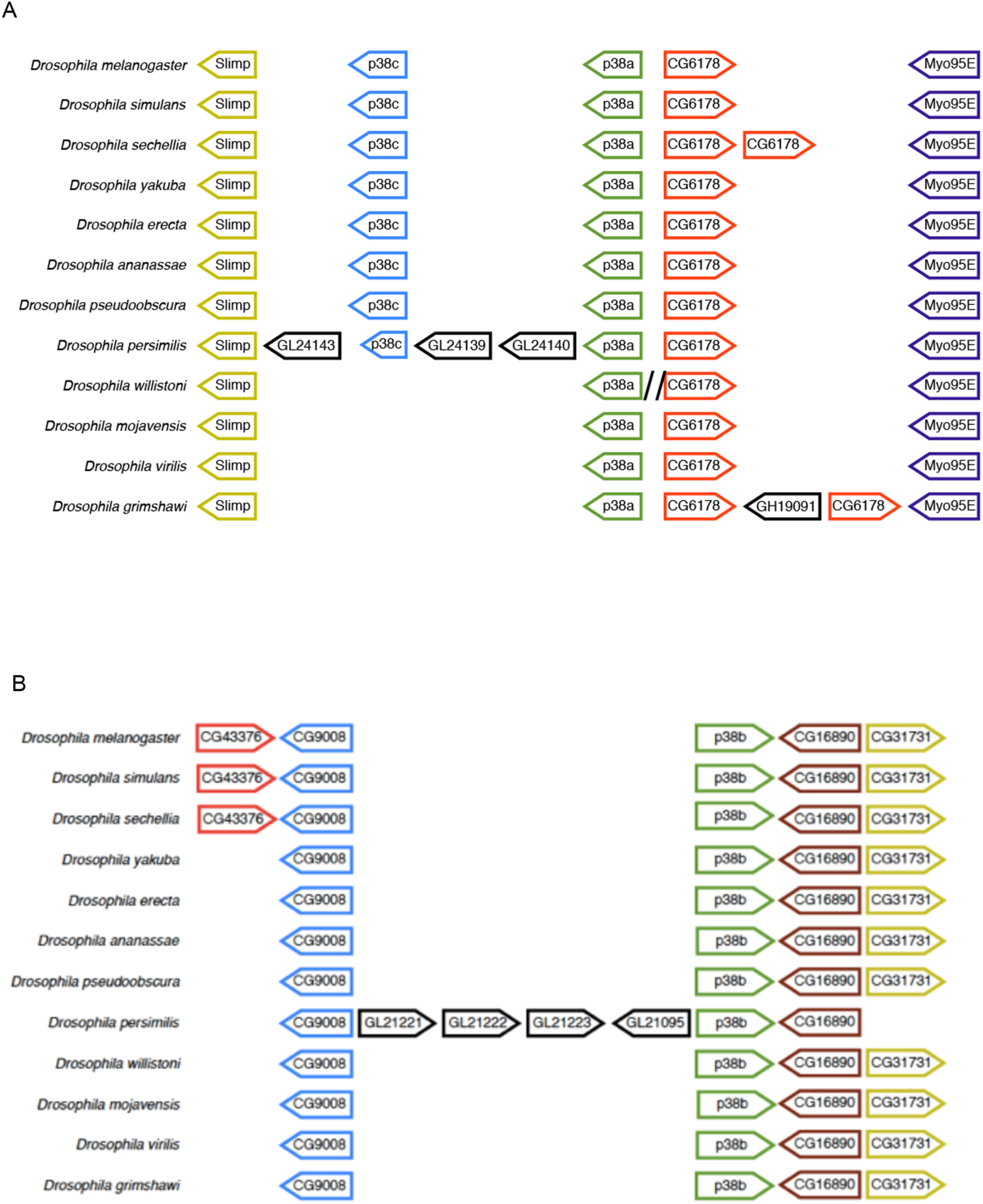
Synteny of the p38K genes in *Drosophila*. A) p38Ka and p38Kc lie in close proximity to each other. *D. persimilis* has additional genes present in this locus as well as a truncated p38Kc gene. *D. sechellia* has a duplication of CG6178 as does *D. grimshawi*, which also has an additional gene inserted into the locus. Backslashes indicate additional inserted DNA with no predicted genes. B) p38Kb synteny is also highly conserved. Within the melanogaster subgroup is the appearance of CG43376. *D. persimilis* has an insertion of 4 genes within this locus and is lacking the CG31731 gene. *D. pseudoobscura* has two p38Kb genes, though the second p38Kb gene is found on a different chromosome.

The synteny of the p38Kb locus is also highly conserved (Figure 3B). The melanogaster subgroup has an additional gene (CG43376) inserted into this region. Furthermore, much like the p38Ka and p38Kc locus, *D. persimilis* has an insertion of four genes (GL21221, GL21222, GL21223, and GL21095) as well as loss of the gene CG31731 in this region (Figure 3B). Though *D. pseudoobscura* has two p38Kb genes (Figure 2C), these genes reside on different chromosomes with p38Kb on the second chromosome (Muller element E) and p38Kb 2 on the fourth chromosome (Muller element B).

### The p38K genes are under purifying selection

Since the p38K genes are conserved across *Drosophila* species, we investigated if these genes show evidence of selection. We analyzed the dN/dS ratio for each of the p38K genes in 22 different species of *Drosophila* (Figure 4). Both p38Ka and p38Kb are under strong purifying selection with average dN/dS ratios of 0.0741 and 0.0479, respectively (Figure 4). This suggests that amino acid substitutions are not well tolerated in these genes and that they have evolved important functions that when disrupted lead to decreased fitness. Interestingly, the p38Ka and p38Kb proteins are also very similar to each other. For example, in *D. melanogaster* the two proteins are 78.4% identical and 94.5% similar at the amino acid level. Therefore, the limited amino acid differences between the p38Ka and p38Kb proteins may be important for their independent functions.

**Figure 4.**
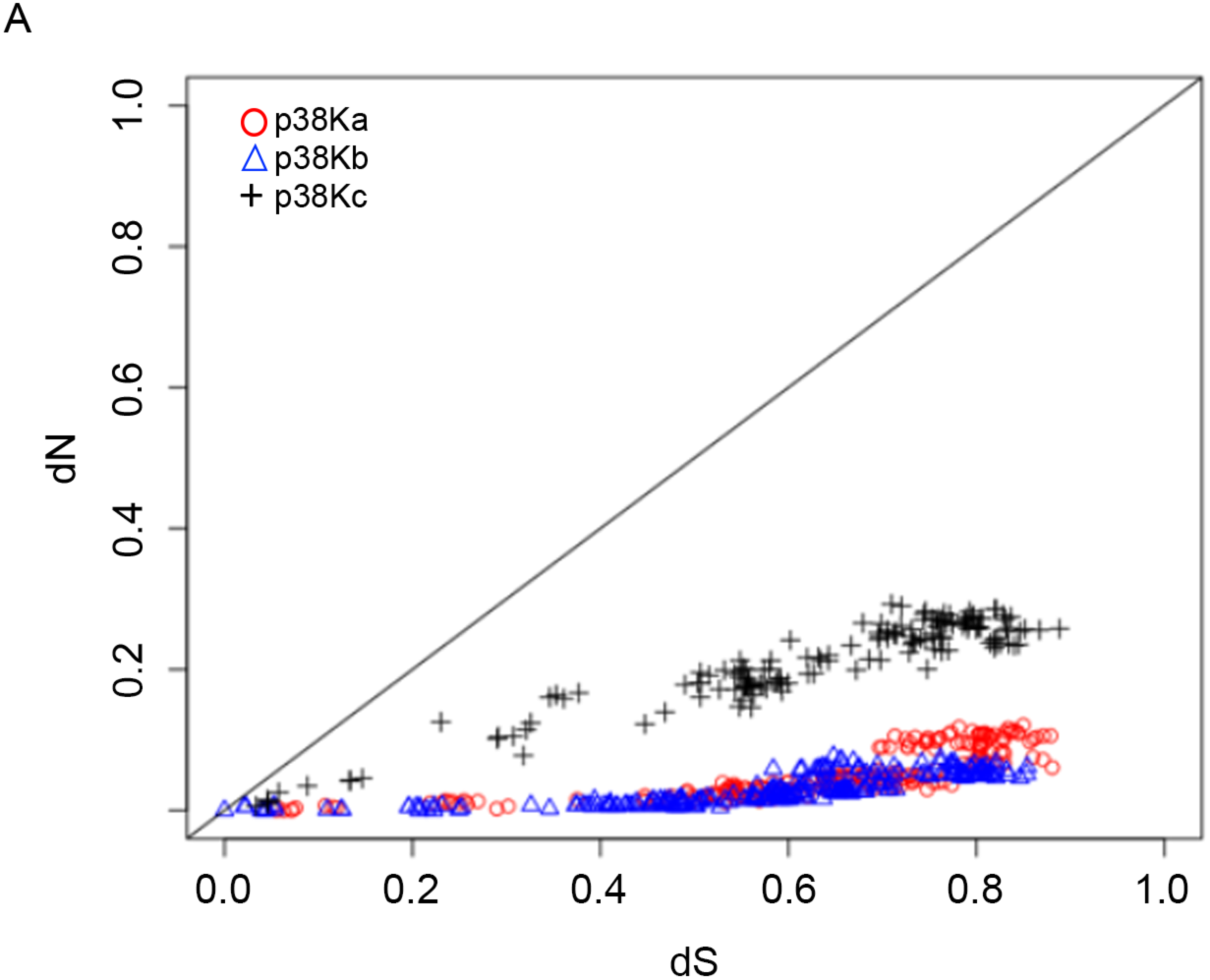
The p38K genes are under purifying selection. Plot of the dN and dS values for the p38K family genes (p38Ka shown in red circles, p38Kb in blue triangles and p38Kc in black crosses). Each spot represents a pairwise comparison between two species.

We also find that p38Kc is under purifying selection but has a higher dN/dS ratio (dN/dS ratio of 0.3346, Figure 4). These data suggest that p38Kc is less constrained than p38Ka and p38Kb, which allowed not only the diversity in amino acid sequence across species, but potentially the evolution of new functions. Furthermore, as p38Kc is under selective pressure, this suggests that these novel functions are also critical for the well-being of the organism.

### Identification of conserved transcription factor binding sites for p38Ka and p38Kb

Though the *D. melanogaster* p38Ka and p38Kb genes are strikingly similar to each other and have some redundant functions (*4–6*), these genes have also been shown to have some independent functions (*4–9,14*). We have also previously reported that the p38Kb protein is highly expressed in muscle and brain, whereas p38Ka is expressed at lower levels in these tissues (*4*), which may contribute to the specific functions of each of these genes. Therefore, we analyzed 1kb upstream of each gene for putative transcription factor binding sites in the 12 *Drosophila* species. We find that p38Ka has four consensus transcription factor binding sites (Figure 5A). Three of these motifs are mostly conserved across *Drosophila* species within the melanogaster group. The first site is a consensus homeobox binding site, which can be bound by members of the Hox family of transcription factors. The other two conserved binding sites are consensus sequences for two isoforms of the transcription factor lola: lola-PO and lola-PK. The final binding site (motif 3) is for an unknown transcription factor and is found only in *D. erecta*.

**Figure 5.**
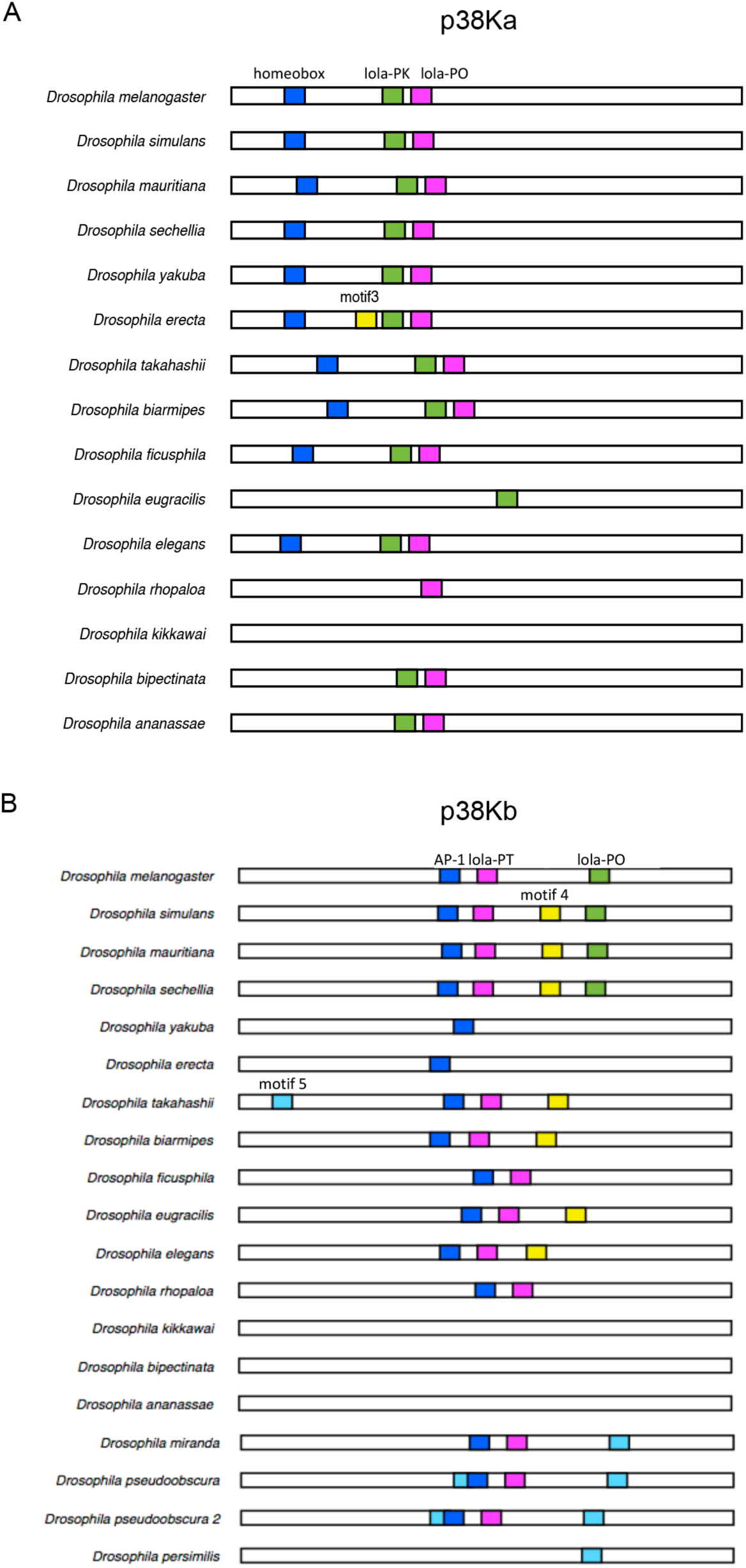
Transcription factor binding sites upstream of p38Ka and p38Kb. The 1Kb upstream region for A) p38Ka and B) p38Kb was analyzed for conserved transcription factor binding sites.

In comparison, p38Kb has five consensus transcription factor binding sites (Figures 5B and S2) that are conserved across the melanogaster and obscura species groups. Similar to p38Ka, p38Kb has two consensus sites for isoforms of the lola transcription factor, lola-PT and lola-PO. p38Kb also has a widely conserved site which corresponds to the AP-1 transcription factor binding site. The final two motifs represent binding sties for unknown transcription factors. Motif 4 is conserved within the obscura group and also in the more distantly related species *D. takahashii*. Motif 5 may be a binding site that has been repeatedly gained or lost over evolutionary history within the melanogaster species group. Interestingly, Motif 5 has been lost in *D. melanogaster* but retained in its closely related sister species *D. simulans, D. mauritiana* and *D. sechellia*. The lola-PO binding site, on the other hand, is restricted to *D. melanogaster* and its sister species. Furthermore, we identified both AP-1 and lola-PT binding sites in most of the species analyzed.

Both the p38Ka and p38Kb loci include binding sites for transcription factors that play a role in development (the Hox family and AP-1 (*25–28*)). Though loss of either p38Ka or p38Kb is viable, double knockout animals are inviable (*4–7,9*), suggesting that p38K signaling may play an important role in developmental processes much like in mammalian systems (*29*). In addition to its role in development, AP-1, a heterodimer of the transcription factors jun and fos, also plays a role in immune functions (*30,31*) and oxidative stress response (*32*). Interestingly, p38Kb has been shown to phosphorylate jun (*33*), suggesting that p38Kb expression may be regulated through a feedback loop.

lola is a key regulator of axon guidance (*34–36*). However, lola has also been shown to act in the ovary to regulate programmed cell death in both developmental and starvation conditions (*37*). Furthermore, the ModENCODE project demonstrated that lola is highly expressed in response to adult exposure to the oxidizing agent paraquat (*38,39*), though the role of specific isoforms of lola in oxidative stress are not well characterized.

### AP-1 and lola-PT regulate the expression of p38Kb in D. melanogaster

Oxidative stress is known to induce the phosphorylation and thus activation of p38K proteins in a variety of systems including *D. melanogaster*. However, the presence of both the AP-1 and lola isoform sites suggests that p38K transcription may also be regulated by oxidative stress. Therefore, we tested if these conserved transcription factor binding sites are important for the regulation of the p38Kb gene. We focused on the AP-1 and lola-PT binding sites since they are exclusive to p38Kb and have links to MAPK signaling pathways and/or oxidative stress. We over-expressed either AP-1 or lola-PT in the muscle using the MHC-GAL4 driver as p38Kb has been shown to act in this tissue to regulate viability and the oxidative stress response (*4*). We find that over-expression of AP-1 is able to induce expression of p38Kb under normal conditions, but not when the flies are exposed to the oxidizing agent paraquat (Figure 6A-B). Over-expression of lola-PT induced p38Kb expression in both normal and oxidative stress conditions (Figure 6C-D). These data suggest that both AP-1 and lola-PT are capable of mediating p38Kb transcription and that lola-PT may be playing a role in the oxidative stress response by promoting the transcription of p38Kb during stress conditions. As transgenic over-expression of p38Kb in the muscle leads to increased lifespan (*4*), we next tested the effect of AP-1 and lola-PT on lifespan. We find that neither over-expression of AP-1 or lola-PT alters lifespan (Figure S3, Table S7, χ^2^=1.8, p-value=0.4 and χ^2^= 5.5, p-value=0.06, respectively), suggesting that the levels of increased p38Kb transcription induced by AP-1 or lola-PT are not sufficient to extend lifespan. Over-expression of p38Kb is also protective against oxidative stress exposure (*4*). Therefore, we assessed the effects of AP-1 and lola-PT over-expression on resistance to paraquat. We find that over-expression of AP-1 leads to increased survival in response to both high (20mM) or low (10mM) paraquat exposure (Figure 6E and G, Tables S8 and S9). Interestingly, lola-PT over-expression also leads to increased survival in response to oxidative stress but only in response to low dose paraquat exposure (Figure 6F and H, Tables S8 and S9). These data suggest that AP-1 and lola-PT are able to induce sufficient levels of p38Kb that leads to a readily available pool of p38K that can be activated and induce the oxidative stress response. However, lola-PT mediated protection is limited to the degree of oxidative stress exposure.

**Figure 6.**
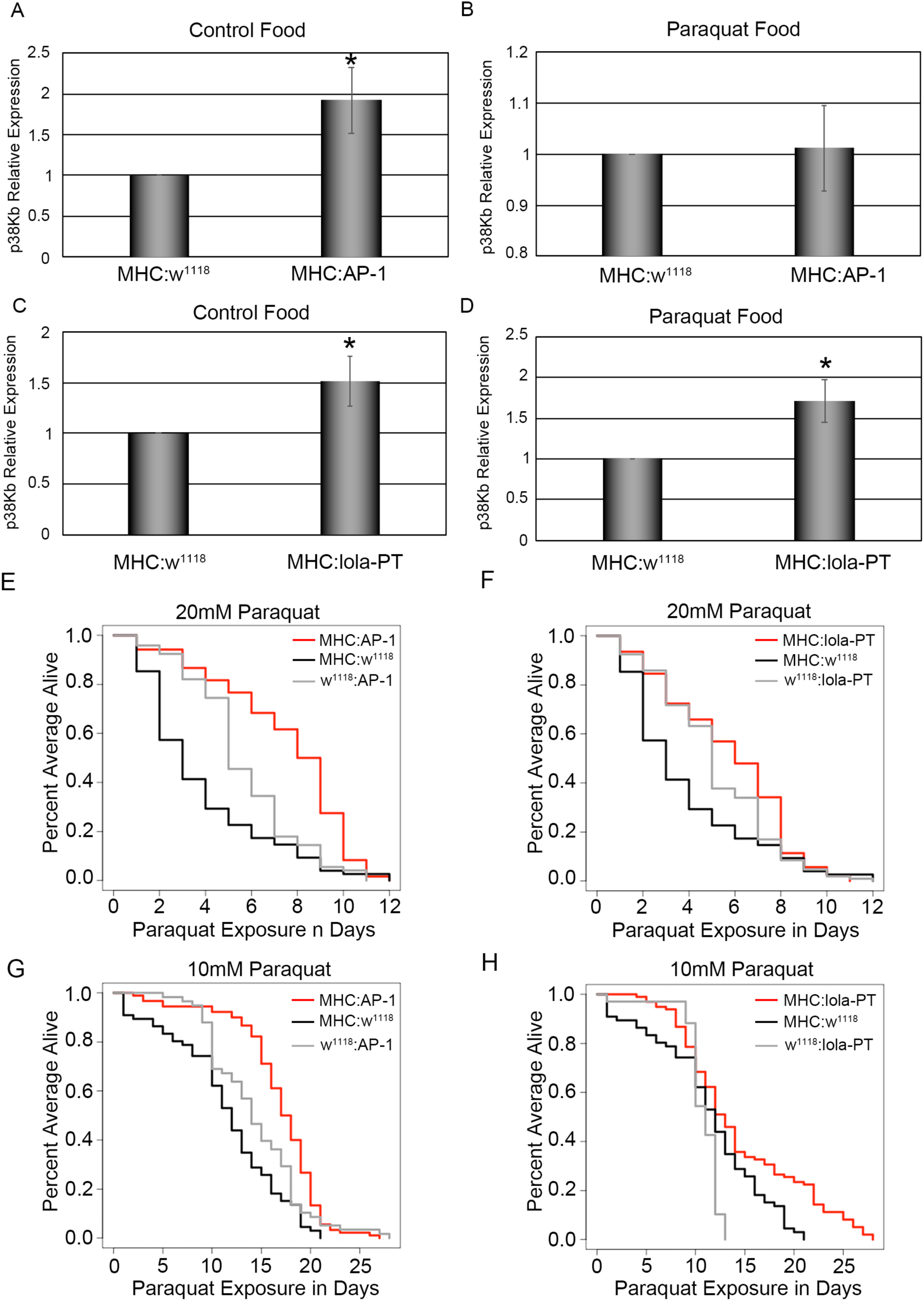
AP-1 and lola-PT mediate p38Kb transcription and oxidative stress resistance. Levels of p38Kb transcription in response to over-expression of AP-1 under A) control conditions and B) 20mM paraquat. Levels of p38Kb transcription in response to over-expression of lola-PT under C) control conditions and D) 20mM paraquat. Resistance to 20mM paraquat was measured for E) AP-1 (red) or F) lola-PT (red) over-expression flies. Resistance to 10mM paraquat G) AP-1 (red) or H) lola-PT (red) over-expression flies. Outcrossed GAL4 and transgene controls are in black and grey, respectively.

### Assessing the relationship between p38K and survival across species

To explore the evolution of oxidative stress response across *Drosophila*, we tested the resistance of 7 species to exposure with the oxidizing agent paraquat. We find that there are significant differences in survival between different species (Figure 7A and Table S10). Interestingly, *D. virilis*, the largest species in our study (*24*) did not have the longest survival time on paraquat (Figure 7A and Table S10). Accordingly, we find that body size and paraquat resistance are only weakly correlated between species (Figure S4A, R=0.44, p-value=0.4529) or when considering phylogenetically independent contrasts (Figure S4B, R=-0.4, p-value=0.6), suggesting that body size is not a major determinant of stress resistance.

**Figure 7.**
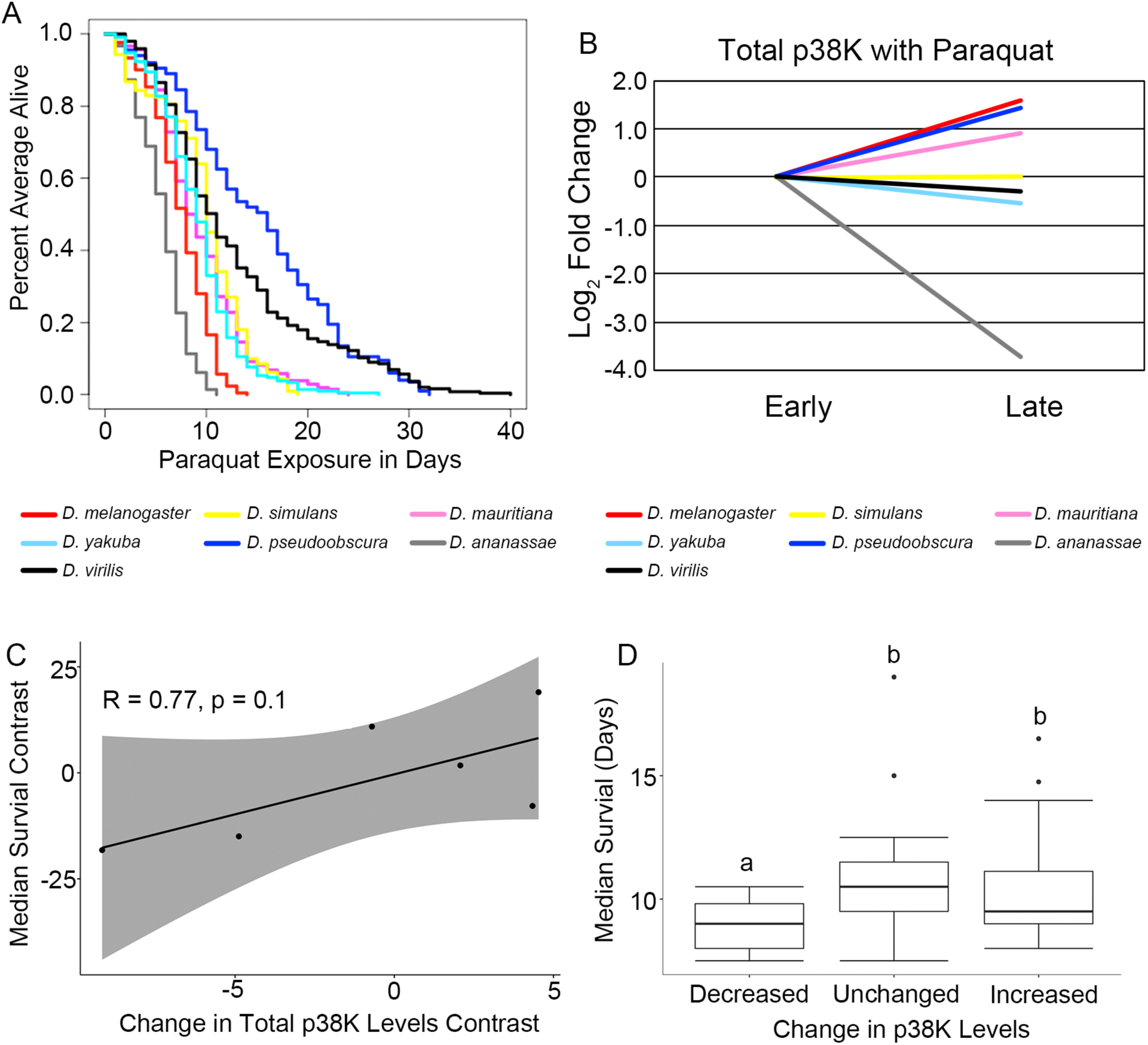
Oxidative stress is correlated with p38K protein levels. A) Kaplan-meier plot of species survival during exposure to 20mM paraquat. B) Log_2_ fold change of levels of p38K protein during paraquat exposure. C) Scatterplot of the correlation between phylogenetically independent contrasts of p38K protein levels and median survival time. D) Median survival of *Drosophila* species with decreased, unchanged or increased levels of p38K during paraquat exposure. a,b indicate significance groups with p < 0.05.

We find that of the studied species, *D. pseudoobscura* showed the longest survival in response to paraquat (Figure 7A and Table S10). This finding is especially interesting as *D. pseudoobscura* has two copies of the p38Kb gene, suggesting that p38Kb may be important for stress resistance and that both copies may contribute to this response. To examine the potential role of p38K, flies were exposed to either control food or food mixed with 20mM paraquat and collected at early and late timepoints based on the average survival time for that species. We performed immunoblots to analyze changes in the levels of p38K protein. We observe a decrease in p38K protein levels on control food with time across the species (Figure S5A; p = 0.026), and we find no significant differences between species (p = 0.40), suggesting that in general, p38K protein levels decline with age in *Drosophila* species.

When flies were subjected to prolonged paraquat exposure, we find an increase in the levels of p38K in *D. melanogaster*, *D. mauritiana*, and *D. pseudoobscura* (log_2_ fold change greater than 0.5), a lack of change in *D. simulans* and *D. virilis* (log_2_ fold change between -0.5 and 0.5), and decreased levels in *D. yakuba* and *D. ananassae* (log_2_ fold change less than -0.5, Figure 7B).

We next tested if the changes in p38K protein levels are associated with species survival when exposed to paraquat. To account for the potential role of phylogenetic relationships in survival, we compared phylogenetically independent contrasts for changes in p38K protein levels and species survival. We find that there is a moderate positive correlation between the change in p38K levels and paraquat resistance (Figure 7C, R = 0.77, p = 0.1). When we categorized the species by changes in total p38K as increased, unchanged, or decreased, we find that decreased levels of p38K are associated with poor survival on paraquat when compared to those species with neutral or increased p38K (Figure 7D, p-value=0.00429).

It has been well documented that oxidative stress induces the activation of p38K through its phosphorylation (*2*). We performed Western blots on our samples using an anti-phosphorylated p38K antibody and find that across species and conditions, there is a linear relationship between the changes in levels of total and phosphorylated forms of p38K (Figure S5B-D).

### Paraquat resistance and p38K protein levels across Drosophila taxa are associated with AP-1 and lola-PT binding sites

Since over-expression of either AP-1 or lola-PT is able to induce p38Kb transcription in *D. melanogaster*, we next tested if having a conserved AP-1 and/or lola-PT site is important for paraquat resistance. Therefore, we tested 7 species of *Drosophila* with different combinations of these sites (Figure 5B) for oxidative stress resistance. Two of these species, *D. simulans* and *D. mauritiana*, are sister species of *D. melanogaster* that have both the AP-1 and lola-PT binding sites as well as a conserved motif not found in *D. melanogaster*. *D. yakuba* is also a member of the melanogaster subgroup and has the AP-1 binding site but not the lola-PT site. *D. pseudoobscura* is a more distantly related species and has two p38Kb genes both of which have AP-1 and lola-PT binding sites. Finally, we tested two species that do not have the conserved AP-1 or lola-PT sites, *D. ananassae* and *D. virilis*, which is the most distantly related of the species. In order to assess the roles of the five predicted transcription factor binding sites, we used ANOVA on Aligned Rank Transformed data to test for interactions between the presence of a transcription factor binding site and p38K protein levels. We first tested the relationship between p38K levels and the presence of lola-PO, motif 4 or motif 5, which are either also present in the p38Ka locus or have no known association with oxidative stress. We find that across species has no significant interaction with any of these three binding sites and p38K protein levels under any conditions (Table S11).

Though p38K levels tend to decrease across time (Figure S5), we find that the presence of an AP-1 site is associated with a smaller decrease in p38K levels under standard conditions (Figure 8A, p=0.037). In addition, the AP-1 site is also associated with increased levels of p38K on paraquat food (Figure 8A, p = 0.037), however, there is no interaction between AP-1 site presence and food treatment (Figure 8A, p = 0.175), suggesting that AP-1 regulation of p38K levels is not responsive to paraquat treatment, in line with our qPCR results (Figure 6A-B). The presence of a lola-PT binding site leads to increased levels of p38K on paraquat but not control food (Figure 8B), and significantly interacts with treatment (Figure 8B, p = 0.013), suggesting that lola-PT is regulating p38K levels in response to oxidative stress. We find that the presence of an AP-1 or lola-PT binding site shows a positive trend toward increasing median survival on paraquat food, although these effects fall just short of significance (Figure 8C-D, p = 0.098 for AP-1, p = 0.069 for lola-PT). These data suggest that AP-1 plays a role in the normal regulation of p38Kb transcription, whereas lola-PT acts in response to oxidative stress to increase p38Kb transcription, and that both modes of regulation can contribute to p38K-mediated oxidative stress resistance.

**Figure 8.**
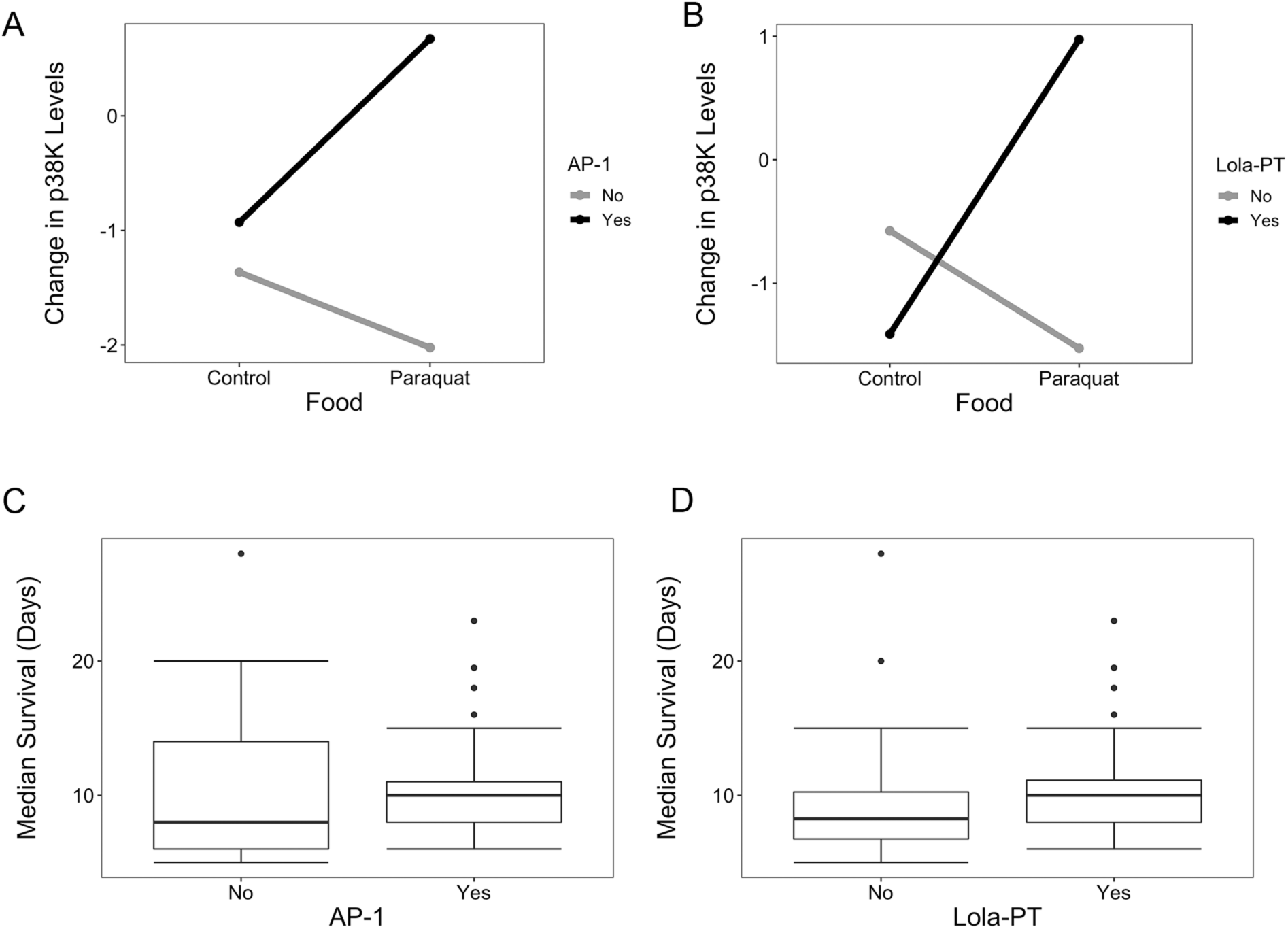
The presence of AP-1 and lola-PT binding sites are associated with p38K protein levels. A) The presence of an AP-1 site is associated with increased levels of p38K protein but does not interact with food treatment. B) The presence of a lola-PT site is associated with increased p38K levels during paraquat treatment and interacts with paraquat treatment . Median survival of *Drosophila* species with C) AP-1 and D) lola-PT sites.

## Discussion

A common theme in biology is the repurposing of signaling pathways to perform new cellular functions. Evolutionary pressures can act on these signaling pathways in various ways leading to sequence divergence, gene duplications or the creation/loss of regulatory elements. In order to understand how these pressures may have influenced an organism’s ability to respond to oxidative stress, we utilized the power of the *Drosophila* genome sequencing projects, which have resulted in the sequencing of 22 *Drosophila* species, 12 of which have been fully annotated. We undertook a comparative genomics approach to analyze the genomic architecture of the conserved p38K genes across the genus *Drosophila* which represents 30-40 million years of evolutionary history (*40*). We find a recent duplication event most likely of p38Ka resulted in formation of the p38Kc gene, which has diverged from both p38Ka and p38Kb and has potentially developed new functions as has been observed in *D. melanogaster* (*4–9,11–16,33,41,42*). In addition, analysis of the 1kb upstream region revealed loss and gain of transcription factor binding sites for both the p38Ka and p38Kb loci. We find that p38Ka and p38Kb both share lola-PO binding sites, which likely plays a role in regulating the expression of both p38Ka and p38Kb in shared functions. Additionally, each gene has a specific set of transcription factor binding sites that may play a role in regulating the expression of these genes for specific functions.

To determine if this bioinformatics approach identified biologically relevant regulatory elements, we tested the function of these sites *in vivo* using the genetically amenable species *D. melanogaster*. We find that the p38Kb transcription factor binding sites for both AP-1 and lola-PT are functional as over-expression of AP-1 and lola-PT induce expression of p38Kb under control conditions. In addition, we also found that these sites play different roles in the regulation of p38Kb as only over-expression of lola-PT was able to further induce expression of p38Kb in the presence of the oxidizing agent paraquat. Intriguingly, we find that over-expression of AP-1 in which flies are aged a week before being exposed to paraquat leads to increased resistance compared to lola-PT over-expression. These data suggest that AP-1 may be able to transcribe sufficient amounts of p38Kb pre-exposure that results in a protective effect once the animal encounters an oxidizing environment. Thus, AP-1 and lola-PT are playing unique roles in the regulation of p38Kb expression and this may, in turn, help the organism as it navigates environmental changes. To determine if these sites may play similar roles in other *Drosophila* species, we tested how species with differing combinations of the AP-1 and lola-PT binding sites respond to oxidative stress. Similar to our results with *D. melanogaster*, we find a significant interaction between the presence of an AP-1 site and increased levels of p38K protein regardless of the oxidative stress state, whereas the presence of a lola-PT binding site shows a significant interaction with p38K protein levels only in the presence of oxidative stress. Furthermore, we found a positive trend between the presence of these sites and increased survival. However, these results were not significant, likely due to the limited number of species identified with different combinations of AP-1 and lola-PT binding sites that we could test. Overall our data suggest that AP-1 and lola-PT act in a similar manner across species to regulate p38Kb expression. Much of what is known about how p38K proteins regulate stress responses has focused on the role of post-translational modifications, in particular, the dual phosphorylation of p38K (*1,3,43*), and little is known about the role of transcriptional regulation in this process. We find that over-expression of either AP-1 or lola-PT is sufficient to increase p38Kb transcription under non-stress conditions and confer resistance to subsequent oxidative stress. These findings suggest that transcriptional regulation of p38Kb can also play an important role in regulating the oxidative stress response, and the transcription of p38Kb in response to oxidative stress may complement the effects of p38Kb phosphorylation in promoting resistance. However, more research will be necessary to understand the relationship between p38Kb induced transcription and subsequent phosphorylation and how this supports the oxidative stress response.

Our results suggest that we can successfully utilize the genomes of sequenced species to identify biologically relevant regulatory elements that may give insights into how different species have evolved mechanisms for responding to selective pressures. Furthermore, as little is known about the transcriptional regulation of other MAP Kinases, our comparative genomics approach may be utilized to identify conserved transcription factor binding sites that may play important roles in regulating these MAP Kinases and potentially other key signaling genes in response to a variety of cellular or environmental stimuli.

**Figure S1. Phylogenetic tree of the p38K genes in *Drosophila*.** The evolutionary relationships between all three p38K genes across 22 species of *Drosophila*. Black denotes the p38Ka genes, red the p38Kb genes, and grey the p38Kc genes.

**Figure S2. Consensus sequences the p38Kb transcription factor binding sites.** The consensus sequence of the A) lola-PT, B) AP-1, C) lola-PO, D) Motif 4, and E) Motif 5 transcription factor binding sites. For each binding site, the size of the letter indicates the proportion of confirmed sites that use that base.

**Figure S3. Over-expression of AP-1 or lola-PT has no effect on lifespan.** A) Over-expression of AP-1 (red) or B) lola-PT (red) reared on control food has no significant effect on lifespan as compared to outcrossed GAL4 (black) and transgene (grey) controls.

**Figure S4. Body size is weakly correlated with paraquat resistance.** A) Scatterplot of correlation between female thorax length and median survival. B) Scatterplot of correlation between phylogenetically independent contrasts of female thorax length and median survival.

**Figure S5. Levels of total and phosphorylated p38K have a linear relationship.** A) Log_2_ fold change of levels of p38K protein on control food. Scatter plots showing the relationship between changes in total and phosphorylated p38K under B) all conditions, C) control conditions and D) exposure to 20mM paraquat.

## Acknowledgements

We would like to acknowledge the Bloomington Drosophila Stock Center (NIH P40OD018537), and the Drosophila Species Stock Center for providing fly stocks used in this study. A. Vrailas-Mortimer was funded by start-up funds from the University of Denver, a Knoebel Center for the Study of Aging pilot grant, start-up funds and a New Faculty Initiative Grant from Illinois State University and was supported by the National Institute of Arthritis and Musculoskeletal and Skin Diseases of the National Institutes of Health under Award Number R15AR070505. The content is solely the responsibility of the authors and does not necessarily represent the official views of the National Institutes of Health. S. Ryan was supported by a Knoebel Center for the Study of Aging pilot grant to A. Vrailas-Mortimer and S. Barbee, N. Mortimer was funded by start-up funds and a Pre-tenure Faculty Initiative Grant from Illinois State University. The funders had no role in study design, data collection and analysis, decision to publish, or preparation of the manuscript.

## References

1. M. Krishna, H. Narang, The complexity of mitogen-activated protein kinases (MAPKs) made simple. Cell. Mol. Life Sci. CMLS. 65, 3525–3544 (2008).

2. J. A. McCubrey, M. M. Lahair, R. A. Franklin, Reactive oxygen species-induced activation of the MAP kinase signaling pathways. Antioxid. Redox Signal. 8, 1775–1789 (2006).

3. W. Peti, R. Page, Molecular basis of MAP kinase regulation. Protein Sci. Publ. Protein Soc. 22, 1698–1710 (2013).

4. A. Vrailas-Mortimer, T. del Rivero, S. Mukherjee, S. Nag, A. Gaitanidis, D. Kadas, C. Consoulas, A. Duttaroy, S. Sanyal, A muscle-specific p38 MAPK/Mef2/MnSOD pathway regulates stress, motor function, and life span in Drosophila. Dev. Cell. 21, 783–795 (2011).

5. J. Chen, C. Xie, L. Tian, L. Hong, X. Wu, J. Han, Participation of the p38 pathway in Drosophila host defense against pathogenic bacteria and fungi. Proc. Natl. Acad. Sci. U. S. A. 107, 20774–20779 (2010).

6. N. Shinzawa, B. Nelson, H. Aonuma, K. Okado, S. Fukumoto, M. Miura, H. Kanuka, p38 MAPK-dependent phagocytic encapsulation confers infection tolerance in Drosophila. Cell Host Microbe. 6, 244–252 (2009).

7. M. Cully, A. Genevet, P. Warne, C. Treins, T. Liu, J. Bastien, B. Baum, N. Tapon, S. J. Leevers, J. Downward, A role for p38 stress-activated protein kinase in regulation of cell growth via TORC1. Mol. Cell. Biol. 30, 481–495 (2010).

8. J.-S. Park, Y.-S. Kim, M.-A. Yoo, The role of p38b MAPK in age-related modulation of intestinal stem cell proliferation and differentiation in Drosophila. Aging. 1, 637–651 (2009).

9. C. R. Craig, J. L. Fink, Y. Yagi, Y. T. Ip, R. L. Cagan, A Drosophila p38 orthologue is required for environmental stress responses. EMBO Rep. 5, 1058–1063 (2004).

10. H. Inoue, M. Tateno, K. Fujimura-Kamada, G. Takaesu, T. Adachi-Yamada, J. Ninomiya-Tsuji, K. Irie, Y. Nishida, K. Matsumoto, A Drosophila MAPKKK, D-MEKK1, mediates stress responses through activation of p38 MAPK. EMBO J. 20, 5421–5430 (2001).

11. E.-M. Ha, K.-A. Lee, Y. Y. Seo, S.-H. Kim, J.-H. Lim, B.-H. Oh, J. Kim, W.-J. Lee, Coordination of multiple dual oxidase-regulatory pathways in responses to commensal and infectious microbes in drosophila gut. Nat. Immunol. 10, 949–957 (2009).

12. C. West, N. Silverman, p38b and JAK-STAT signaling protect against Invertebrate iridescent virus 6 infection in Drosophila. PLoS Pathog. 14, e1007020 (2018).

13. J. Na, L. P. Musselman, J. Pendse, T. J. Baranski, R. Bodmer, K. Ocorr, R. Cagan, A Drosophila model of high sugar diet-induced cardiomyopathy. PLoS Genet. 9, e1003175 (2013).

14. A. D. Vrailas-Mortimer, S. M. Ryan, M. J. Avey, N. T. Mortimer, H. Dowse, S. Sanyal, p38 MAP kinase regulates circadian rhythms in Drosophila. J. Biol. Rhythms. 29, 411–426 (2014).

15. M. M. Davis, D. A. Primrose, R. B. Hodgetts, A member of the p38 mitogen-activated protein kinase family is responsible for transcriptional induction of Dopa decarboxylase in the epidermis of Drosophila melanogaster during the innate immune response. Mol. Cell. Biol. 28, 4883–4895 (2008).

16. S. Chakrabarti, M. Poidevin, B. Lemaitre, The Drosophila MAPK p38c regulates oxidative stress and lipid homeostasis in the intestine. PLoS Genet. 10, e1004659 (2014).

17. K. Tamura, D. Peterson, N. Peterson, G. Stecher, M. Nei, S. Kumar, MEGA5: Molecular Evolutionary Genetics Analysis Using Maximum Likelihood, Evolutionary Distance, and Maximum Parsimony Methods. Mol. Biol. Evol. 28, 2731–2739 (2011).

18. D. T. Jones, W. R. Taylor, J. M. Thornton, The rapid generation of mutation data matrices from protein sequences. Bioinformatics. 8, 275–282 (1992).

19. J. Felsenstein, Confidence Limits on Phylogenies: An Approach Using the Bootstrap. Evolution. 39, 783 (1985).

20. K. Tamura, M. Nei, Estimation of the number of nucleotide substitutions in the control region of mitochondrial DNA in humans and chimpanzees. Mol. Biol. Evol. 10, 512–526 (1993).

21. A. L. Franciscovich, A. D. V. Mortimer, A. A. Freeman, J. Gu, S. Sanyal, Overexpression screen in Drosophila identifies neuronal roles of GSK-3 beta/shaggy as a regulator of AP-1-dependent developmental plasticity. Genetics. 180, 2057–2071 (2008).

22. M. Piipari, T. A. Down, H. Saini, A. Enright, T. J. P. Hubbard, iMotifs: an integrated sequence motif visualization and analysis environment. Bioinformatics. 26, 843–844 (2010).

23. S. Gupta, J. A. Stamatoyannopoulos, T. L. Bailey, W. S. Noble, Quantifying similarity between motifs. Genome Biol. 8, R24 (2007).

24. T. A. Markow, P. M. O’Grady, Drosophila biology in the genomic age. Genetics. 177, 1269–1276 (2007).

25. K. K. Perkins, A. Admon, N. Patel, R. Tjian, The Drosophila Fos-related AP-1 protein is a developmentally regulated transcription factor. Genes Dev. 4, 822–834 (1990).

26. J. R. Riesgo-Escovar, E. Hafen, Common and distinct roles of DFos and DJun during Drosophila development. Science. 278, 669–672 (1997).

27. L. Bataillé, J.-L. Frendo, A. Vincent, Hox control of Drosophila larval anatomy; The Alary and Thoracic Alary-Related Muscles. Mech. Dev. 138 **Pt 2**, 170–176 (2015).

28. J. Enriquez, H. Boukhatmi, L. Dubois, A. A. Philippakis, M. L. Bulyk, A. M. Michelson, M. Crozatier, A. Vincent, Multi-step control of muscle diversity by Hox proteins in the Drosophila embryo. Dev. Camb. Engl. 137, 457–466 (2010).

29. A. Keren, Y. Tamir, E. Bengal, The p38 MAPK signaling pathway: a major regulator of skeletal muscle development. Mol. Cell. Endocrinol. 252, 224–230 (2006).

30. T. Kim, J. Yoon, H. Cho, W.-B. Lee, J. Kim, Y.-H. Song, S. N. Kim, J. H. Yoon, J. Kim-Ha, Y.-J. Kim, Downregulation of lipopolysaccharide response in Drosophila by negative crosstalk between the AP1 and NF-kappaB signaling modules. Nat. Immunol. 6, 211–218 (2005).

31. T. Tokusumi, R. P. Sorrentino, M. Russell, R. Ferrarese, S. Govind, R. A. Schulz, Characterization of a lamellocyte transcriptional enhancer located within the misshapen gene of Drosophila melanogaster. PloS One. 4, e6429 (2009).

32. V. J. Milton, H. E. Jarrett, K. Gowers, S. Chalak, L. Briggs, I. M. Robinson, S. T. Sweeney, Oxidative stress induces overgrowth of the Drosophila neuromuscular junction. Proc. Natl. Acad. Sci. U. S. A. 108, 17521–17526 (2011).

33. Y. Sano, H. Akimaru, T. Okamura, T. Nagao, M. Okada, S. Ishii, Drosophila activating transcription factor-2 is involved in stress response via activation by p38, but not c-Jun NH(2)-terminal kinase. Mol. Biol. Cell. 16, 2934–2946 (2005).

34. E. Giniger, K. Tietje, L. Y. Jan, Y. N. Jan, lola encodes a putative transcription factor required for axon growth and guidance in Drosophila. Dev. Camb. Engl. 120, 1385–1398 (1994).

35. K. Madden, D. Crowner, E. Giniger, LOLA has the properties of a master regulator of axon-target interaction for SNb motor axons of Drosophila. Dev. Biol. 213, 301–313 (1999).

36. D. Crowner, K. Madden, S. Goeke, E. Giniger, Lola regulates midline crossing of CNS axons in Drosophila. Dev. Camb. Engl. 129, 1317–1325 (2002).

37. B. P. Bass, K. Cullen, K. McCall, The axon guidance gene lola is required for programmed cell death in the Drosophila ovary. Dev. Biol. 304, 771–785 (2007).

38. J. B. Brown, N. Boley, R. Eisman, G. E. May, M. H. Stoiber, M. O. Duff, B. W. Booth, J. Wen, S. Park, A. M. Suzuki, K. H. Wan, C. Yu, D. Zhang, J. W. Carlson, L. Cherbas, B. D. Eads, D. Miller, K. Mockaitis, J. Roberts, C. A. Davis, E. Frise, A. S. Hammonds, S. Olson, S. Shenker, D. Sturgill, A. A. Samsonova, R. Weiszmann, G. Robinson, J. Hernandez, J. Andrews, P. J. Bickel, P. Carninci, P. Cherbas, T. R. Gingeras, R. A. Hoskins, T. C. Kaufman, E. C. Lai, B. Oliver, N. Perrimon, B. R. Graveley, S. E. Celniker, Diversity and dynamics of the Drosophila transcriptome. Nature. 512, 393–399 (2014).

39. S. E. Celniker, L. A. L. Dillon, M. B. Gerstein, K. C. Gunsalus, S. Henikoff, G. H. Karpen, M. Kellis, E. C. Lai, J. D. Lieb, D. M. MacAlpine, G. Micklem, F. Piano, M. Snyder, L. Stein, K. P. White, R. H. Waterston, modENCODE Consortium, Unlocking the secrets of the genome. Nature. 459, 927–930 (2009).

40. D. J. Obbard, J. Maclennan, K.-W. Kim, A. Rambaut, P. M. O’Grady, F. M. Jiggins, Estimating Divergence Dates and Substitution Rates in the Drosophila Phylogeny. Mol. Biol. Evol. 29, 3459–3473 (2012).

41. A. Vrailas-Mortimer, R. Gomez, H. Dowse, S. Sanyal, A survey of the protective effects of some commercially available antioxidant supplements in genetically and chemically induced models of oxidative stress in Drosophila melanogaster. Exp. Gerontol. 47, 712–722 (2012).

42. V. E. Belozerov, S. Ratkovic, H. McNeill, A. J. Hilliker, J. C. McDermott, In vivo interaction proteomics reveal a novel p38 mitogen-activated protein kinase/Rack1 pathway regulating proteostasis in Drosophila muscle. Mol. Cell. Biol. 34, 474–484 (2014).

43. L. Chang, M. Karin, Mammalian MAP kinase signalling cascades. Nature. 410, 37–40 (2001).

