## Supplemental Figures and Tables for "Evolutionarily Conserved Transcription Factors Drive the Oxidative Stress Response in *Drosophila*"

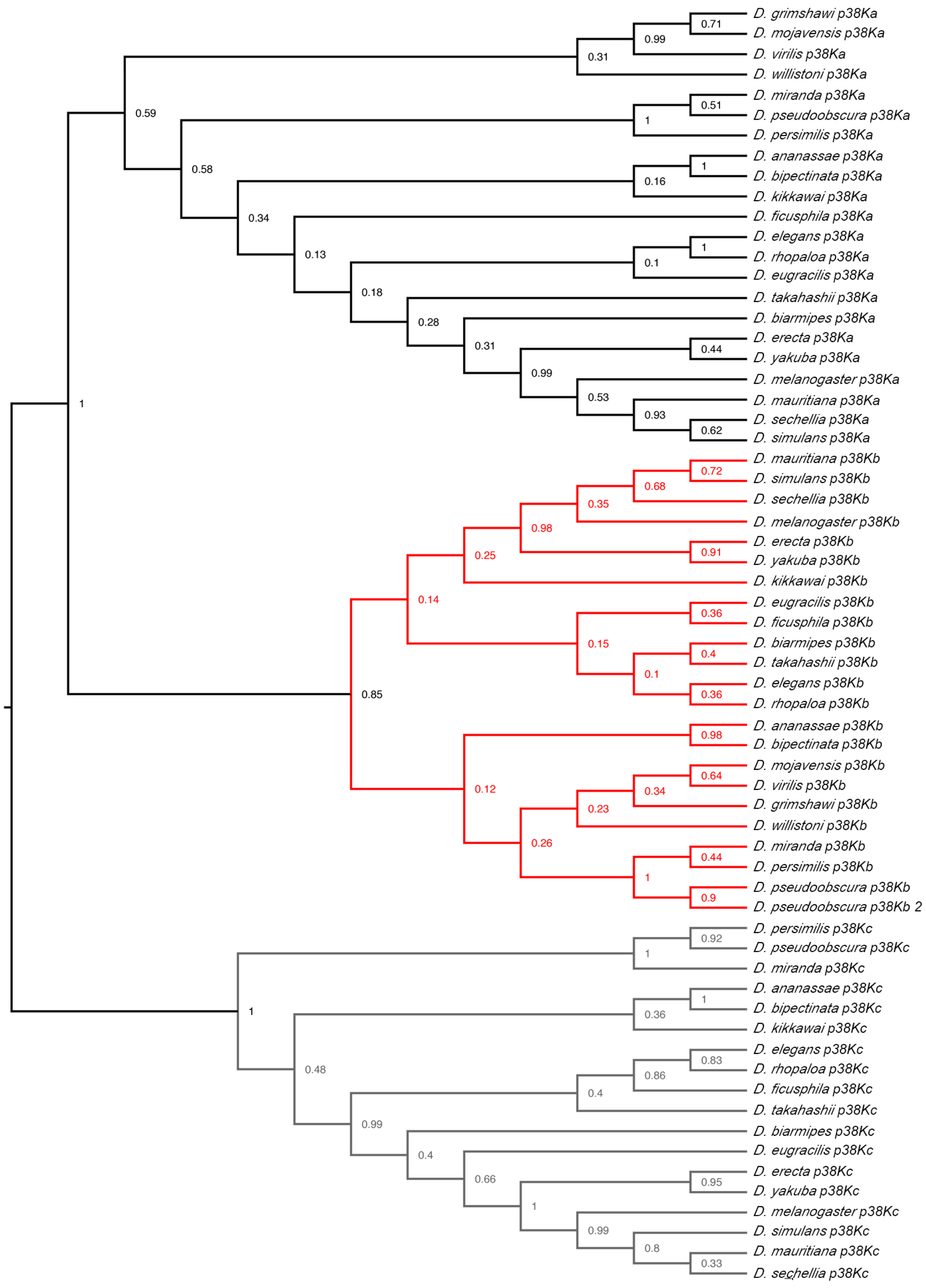
D. grimshawi p38Ka
D. mojavensis p38Ka
D. virilis p38Ka
D. willistoni p38Ka
D. miranda p38Ka
D. pseudoobscura p38Ka
D. persimilis p38Ka
D. ananassae p38Ka
D. bipectinata p38Ka
D. kikkawai p38Ka
D. ficusphila p38Ka
D. elegans p38Ka
D. rhopaloa p38Ka
D. eugracilis p38Ka
D. takahashii p38Ka
D. biarmipes p38Ka
D. erecta p38Ka
D. yakuba p38Ka
D. melanogaster p38Ka
D. mauritiana p38Ka
D. sechellia p38Ka
D. simulans p38Ka
D. mauritiana p38Kb
D. simulans p38Kb
D. sechellia p38Kb
D. melanogaster p38Kb
D. erecta p38Kb
D. yakuba p38Kb
D. kikkawai p38Kb
D. eugracilis p38Kb
D. ficusphila p38Kb
D. biarmipes p38Kb
D. takahashii p38Kb
D. elegans p38Kb
D. rhopaloa p38Kb
D. ananassae p38Kb
D. bipectinata p38Kb
D. mojavensis p38Kb
D. virilis p38Kb
D. grimshawi p38Kb
D. willistoni p38Kb
D. miranda p38Kb
D. persimilis p38Kb
D. pseudoobscura p38Kb
D. pseudoobscura p38Kb 2
D. persimilis p38Kc
D. pseudoobscura p38Kc
D. miranda p38Kc
D. ananassae p38Kc
D. bipectinata p38Kc
D. kikkawai p38Kc
D. elegans p38Kc
D. rhopaloa p38Kc
D. ficusphila p38Kc
D. takahashii p38Kc
D. biarmipes p38Kc
D. eugracilis p38Kc
D. erecta p38Kc
D. yakuba p38Kc
D. melanogaster p38Kc
D. simulans p38Kc
D. mauritiana p38Kc
D. sechellia p38Kc

Figure S1

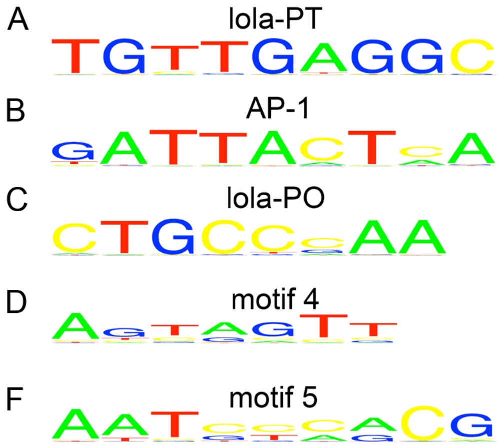


Figure S2

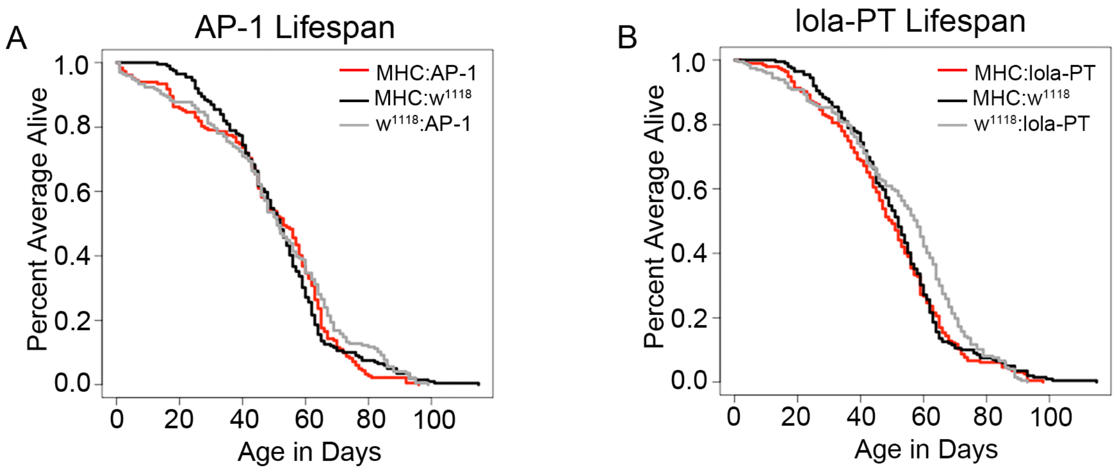

A
AP-1 Lifespan
MHC:AP-1
MHC:w1118
w1118:AP-1
Percent Average Alive
1.0
0.8
0.6
0.4
0.2
0.0
0
20
40
60
80
100
Age in Days
B
lola-PT Lifespan
MHC:lola-PT
MHC:w1118
w1118:lola-PT
Percent Average Alive
1.0
0.8
0.6
0.4
0.2
0.0
0
20
40
60
80
100
Age in Days


Figure S3

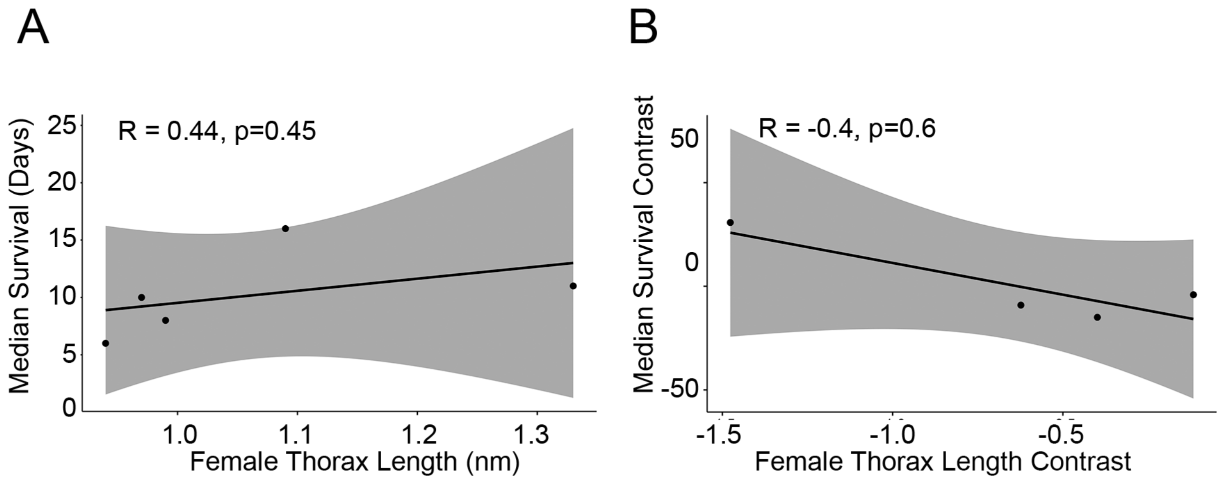
A
R = 0.44, p=0.45
Median Survival (Days)
25
20
15
10
5
0
1.0
1.1
1.2
1.3
Female Thorax Length (nm)
B
R = -0.4, p=0.6
Median Survival Contrast
50
0
-50
-1.5
-1.0
-0.5
Female Thorax Length Contrast

Figure S4

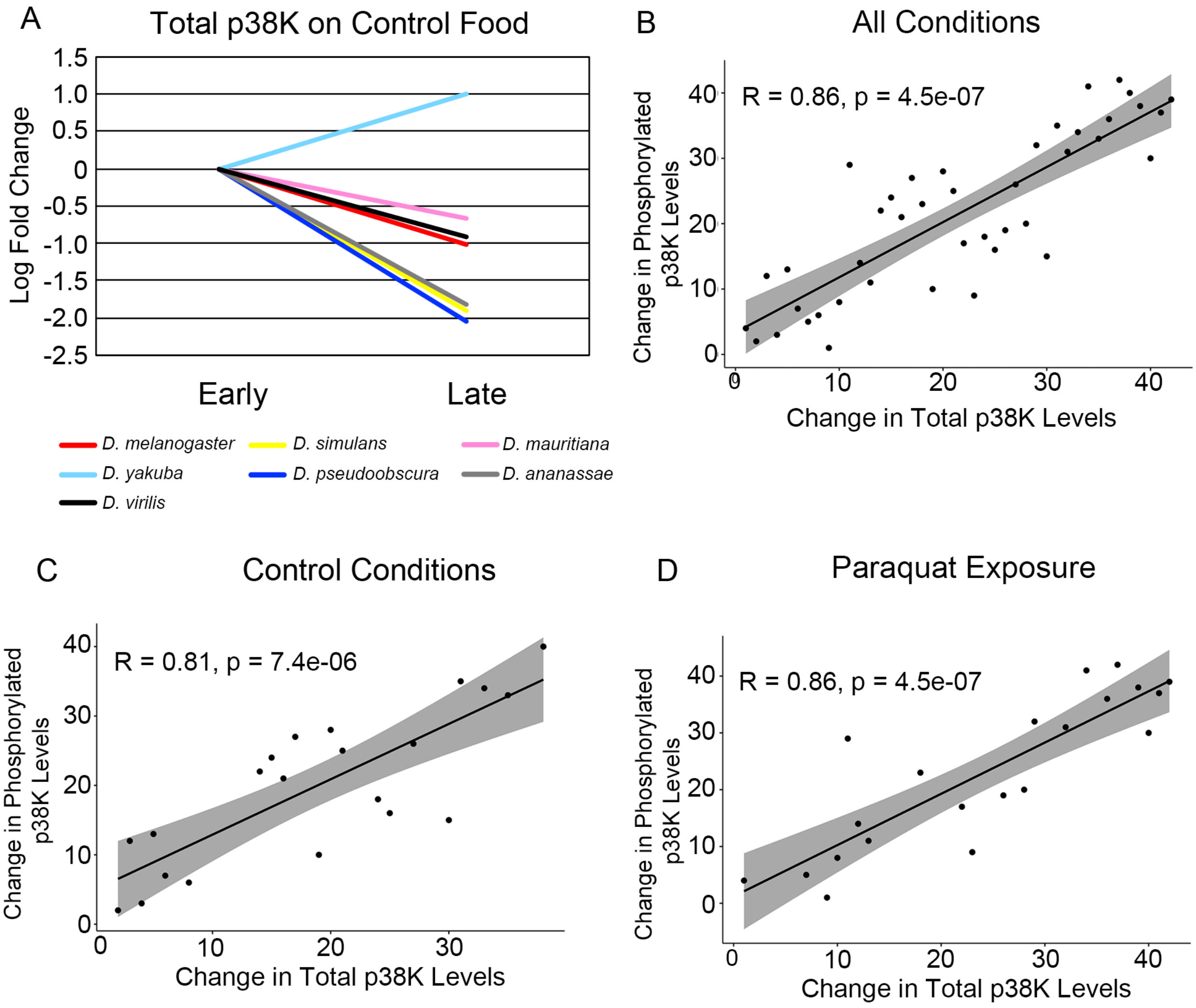

A
Total p38K on Control Food
Log Fold Change
1.5
1.0
0.5
0
-0.5
-1.0
-1.5
-2.0
-2.5
Early
Late
D. melanogaster
D. simulans
D. mauritiana
D. yakuba
D. pseudoobscura
D. ananassae
D. virilis
B
All Conditions
Change in Phosphorylated p38K Levels
R = 0.86, p = 4.5e-07
Change in Total p38K Levels
C
Control Conditions
R = 0.81, p = 7.4e-06
Change in Phosphorylated p38K Levels
Change in Total p38K Levels
D
Paraquat Exposure
R = 0.86, p = 4.5e-07
Change in Phosphorylated p38K Levels
Change in Total p38K Levels


Figure S5

| **Species** | **Gene Name** | **NCBI Accession #** |
|---|---|---|
| *D. ananassae* | p38Ka | XM_001955664.2 |
| *D. biarmipes* | p38Ka | XM_017097085.1 |
| *D. bipectinata* | p38Ka | XM_017249637.1 |
| *D. elegans* | p38Ka | XM_017258256.1 |
| *D. erecta* | p38Ka | XM_001982000.3 |
| *D. eugracilis* | p38Ka | XM_017228726.1 |
| *D. ficusphila* | p38Ka | XM_017191561.1 |
| *D. grimshawi* | p38Ka | XM_001989942.1 |
| *D. kikkawai* | p38Ka | XM_017180763.1 |
| *D. mauritiana* | p38Ka | NIGA01000004.1 (Genome accession ID) |
| *D. melanogaster* | p38Ka | NM_057815.5 |
| *D. miranda* | p38Ka | XM_017289227.1 |
| *D. mojavensis* | p38Ka | XM_001998772.2 |
| *D. persimilis* | p38Ka | XM_002013400.2 |
| *D. pseudoobscura* | p38Ka | XM_001358779.4 |
| *D. rhopaloa* | p38Ka | XM_017125963.1 |
| *D. sechellia* | p38Ka | XM_002032419.1 |
| *D. simulans* | p38Ka | XM_016174818.1 |
| *D. takahashii* | p38Ka | XM_017160399.1 |
| *D. virilis* | p38Ka | XM_002058599.2 |
| *D. willistoni* | p38Ka | XM_002070167.3 |
| *D. yakuba* | p38Ka | XM_002099248.2 |
| *D. ananassae* | p38Kb | XM_001964952.2 |
| *D. biarmipes* | p38Kb | XM_017106505.1 |
| *D. bipectinata* | p38Kb | XM_017250102.1 |
| *D. elegans* | p38Kb | XM_017276942.1 |
| *D. erecta* | p38Kb | XM_001969550.3 |
| *D. eugracilis* | p38Kb | XM_017212644.1 |
| *D. ficusphila* | p38Kb | XM_017192371.1 |
| *D. grimshawi* | p38Kb | XM_001988300.1 |
| *D. kikkawai* | p38Kb | XM_017181846.1 |
| *D. mauritiana* | p38Kb | NIGA01000001.1 (Genome accession ID) |
| *D. melanogaster* | p38Kb | NM_058013.5 |
| *D. miranda* | p38Kb | XM_017296766.1 |
| *D. mojavensis* | p38Kb | XM_002004004.2 |
| *D. persimilis* | p38Kb | XM_026987565.1 |

| | | |
|---|---|---|
| *D. pseudoobscura* | p38Kb | XM_001356214.3 |
| *D. rhopaloa* | p38Kb | XM_017134148.1 |
| *D. sechellia* | p38Kb | XM_002035711.1 |
| *D. simulans* | p38Kb | XM_016180114.1 |
| *D. takahashii* | p38Kb | XM_017159206.1 |
| *D. virilis* | p38Kb | XM_002057421.2 |
| *D. willistoni* | p38Kb | XM_002066223.3 |
| *D. yakuba* | p38Kb | XM_002088597.2 |
| *D. pseudoobscura* | p38Kb2 | XM_002132725.1 |
| *D. ananassae* | p38Kc | XM_001955665.2 |
| *D. biarmipes* | p38Kc | XM_017096660.1 |
| *D. bipectinata* | p38Kc | XM_017249656.1 |
| *D. elegans* | p38Kc | XM_017258257.1 |
| *D. erecta* | p38Kc | XM_026983955.1 |
| *D. eugracilis* | p38Kc | XM_017228501.1 |
| *D. ficusphila* | p38Kc | XM_017191241.1 |
| *D. kikkawai* | p38Kc | XM_017180769.1 |
| *D. mauritiana* | p38Kc | NIGA01000004.1 (Genome accession ID) |
| *D. melanogaster* | p38Kc | NM_206554.2 |
| *D. miranda* | p38Kc | XM_017289226.1 |
| *D. persimilis* | p38Kc | XM_002013401.2 |
| *D. pseudoobscura* | p38Kc | XM_002137454.2 |
| *D. rhopaloa* | p38Kc | XM_017125964.1 |
| *D. sechellia* | p38Kc | XM_002032418.1 |
| *D. simulans* | p38Kc | XM_016174820.1 |
| *D. takahashii* | p38Kc | XM_017160398.1 |
| *D. yakuba* | p38Kc | XM_002099249.2 |

Table S1. Accession numbers for analyzed DNA sequences.

| Species | Gene Name | NCBI Accession # |
|---|---|---|
| *H. sapiens* | ERK1 (MAPK 3) | NP_002737.2 |
| *H. sapiens* | ERK2 (MAPK 1) | NP_002736.3 |
| *M. musculus* | ERK1 (MAPK 3) | NP_036082.1 |
| *M. musculus* | ERK2 MAPK 1 | NP_001033752.1 |
| *D. rerio* | ERK1 (MAPK 3) | NP_958915.1 |
| *D. rerio* | ERK2 (MAPK 1) | NP_878308.2 |
| *D. melanogaster* | rolled | NP_001015122.1 |
| *C. elegans* | mpk-1 | NP_001022584.1 |
| *S. cerevisiae* | FUS3 | NP_009537.1 |
| *S. cerevisiae* | KSS1 | NP_011554.3 |
| *H. sapiens* | JNK1 (MAPK 8) | NP_001310231.1 |
| *H. sapiens* | JNK2 (MAPK 9) | NP_001351536.1 |
| *H. sapiens* | JNK3 (MAPK 10) | NP_001304998.1 |
| *M. musculus* | JNK1 (MAPK 8) | NP_057909.1 |
| *M. musculus* | JNK2 (MAPK 9) | NP_001157144.1 |
| *M. musculus* | JNK3 (MAPK 10) | NP_001075036.1 |
| *D. rerio* | JNK1 (MAPK 8) | NP_571796.1 |
| *D. rerio* | JNK2 (MAPK 9) | XP_001919688.1 |
| *D. rerio* | JNK3 (MAPK 10) | NP_001305247.1 |
| *D. melanogaster* | basket | NP_001162930.1 |
| *C. elegans* | jnk-1 | NP_001021270.1 |
| *S. cerevisiae* | HOG1 | NP_013214.1 |
| *H. sapiens* | p38Ka (MAPK14) | NP_001306.1 |
| *H. sapiens* | p38Kb (MAPK11) | NP_002742.3 |
| *H. sapiens* | p38Kg (MAPK12) | NP_002960.2 |
| *H. sapiens* | p38Kd (MAPK13) | NP_002745.1 |
| *M. musculus* | p38Ka (MAPK14) | NP_036081.1 |
| *M. musculus* | p38Kb (MAPK11) | NP_035291.4 |
| *M. musculus* | p38Kg (MAPK12) | NP_038899.1 |
| *M. musculus* | p38Kd (MAPK13) | NP_036080.2 |
| *D. rerio* | p38Ka (MAPK14) | NP_571797.1 |
| *D. rerio* | p38Kb (MAPK11) | NP_001002095.1 |
| *D. rerio* | p38Kg (MAPK12) | NP_571482.1 |
| *D. rerio* | p38Kd (MAPK13) | XP_001337833.2 |
| *D. melanogaster* | p38Ka | NP_001163711.1 |
| *D. melanogaster* | p38Kb | NP_477361.1 |
| *D. melanogaster* | p38Kc | NP_996277.1 |

| | | |
|---|---|---|
| *C. elegans* | pmk-1 | NP_501365.1 |
| *C. elegans* | pmk-2 | NP_741457.2 |
| *C. elegans* | pmk-3 | NP_501363.1 |
| *D. ananassae* | basket | XP_001963064.2 |
| *D. ananassae* | rolled | XP_001965786.2 |
| *D. virilis* | basket | XP_002052403.1 |
| *D. virilis* | rolled | XP_002052382.1 |
| *D. yakuba* | basket | XP_002089054.1 |
| *D. yakuba* | rolled | XP_002086048.2 |
| *D. ananassae* | p38Ka | XP_001955700.1 |
| *D. ananassae* | p38Kb | XP_001964988.1 |
| *D. ananassae* | p38Kc | XP_001955701.1 |
| *D. virilis* | p38Ka | XP_002058635.2 |
| *D. virilis* | p38Kb | XP_002057457.1 |
| *D. yakuba* | p38Ka | XP_002099284.1 |
| *D. yakuba* | p38Kb | XP_002088633.1 |
| *D. yakuba* | p38Kc | XP_002099285.1 |

Table S2. Accession numbers for analyzed protein sequences.

| Genotype | Average Age batch 1 | Average Age batch 2 | Median Age batch 1 | Median Age batch 2 |
|---|---|---|---|---|
| MHC-GAL4::w1118 | 8.2 days | 3.8 days | 9 days | 3 days |
| MHC-GAL4::AP-1 | 10.3 days | 7.5 days | 10 days | 8.5 days |
| MHC-GAL4::lola PT | 9.3 days | 5.7 days | 10 days | 6 days |

Table S3. Comparisons of different batches of 20mM paraquat.

| Species | Gene Name |
| --- | --- |
| *H. sapiens* | ERK1 (MAPK 3) |
| *H. sapiens* | ERK2 (MAPK 1) |
| *M. musculus* | ERK1 (MAPK 3) |
| *M. musculus* | ERK2 (MAPK 1) |
| *D. rerio* | ERK1 (MAPK 3) |
| *D. rerio* | ERK2 (MAPK 1) |
| *D. melanogaster* | rolled |
| *C. elegans* | mpk-1 |
| *S. cerevisiae* | FUS3 |
| *S. cerevisiae* | KSS1 |

Table S4. ERK MAPK genes across taxa.

| Species | Gene Name |
|---|---|
| *H. sapiens* | JNK1 (MAPK 8) |
| *H. sapiens* | JNK2 (MAPK 9) |
| *H. sapiens* | JNK3 (MAPK 10) |
| *M. musculus* | JNK1 (MAPK 8) |
| *M. musculus* | JNK2 (MAPK 9) |
| *M. musculus* | JNK3 (MAPK 10) |
| *D. rerio* | JNK1 (MAPK 8) |
| *D. rerio* | JNK2 (MAPK 9) |
| *D. rerio* | JNK3 (MAPK 10) |
| *D. melanogaster* | basket |
| *C. elegans* | jnk-1 |
| *S. cerevisiae* | HOG1 |

Table S5. JNK MAPK genes across taxa.

| Species | Gene Name |
|---|---|
| *H. sapiens* | p38K α (MAPK14) |
| *H. sapiens* | p38K β (MAPK11) |
| *H. sapiens* | p38K γ (MAPK12) |
| *H. sapiens* | p38K δ (MAPK13) |
| *M. musculus* | p38K α (MAPK14) |
| *M. musculus* | p38K β (MAPK11) |
| *M. musculus* | p38K γ (MAPK12) |
| *M. musculus* | p38K δ (MAPK13) |
| *D. rerio* | p38K α (MAPK14) |
| *D. rerio* | p38K β (MAPK11) |
| *D. rerio* | p38K γ (MAPK12) |
| *D. rerio* | p38K δ (MAPK13) |
| *D. melanogaster* | p38Ka |
| *D. melanogaster* | p38Kb |
| *D. melanogaster* | p38Kc |
| *C. elegans* | pmk-1 |
| *C. elegans* | pmk-2 |
| *C. elegans* | pmk-3 |
| *S. cerevisiae* | HOG1 |

Table S6. p38K MAPK genes across taxa.

| Genotype | Average Age | Median Age | n | p value vs outcrossed GAL4 | p value vs outcrossed transgene |
|---|---|---|---|---|---|
| MHC-GAL4::w1118 | 51.5 days | 52 days | 199 | n/a | n/a |
| w1118::AP-1 | 50.5 days | 51 days | 196 | 0.54 | n/a |
| MHC-GAL4::AP-1 | 49 days | 53 days | 182 | 0.98 | 0.39 |
| w1118::lola PT | 53.0 days | 58 days | 197 | 0.154 | n/a |
| MHC-GAL4::lola PT | 49.1 days | 50 days | 195 | 0.472 | 0.099 |

Table S7. Lifespan of AP-1 and lola-PT over-expression. $\chi2=1.8$, $p=0.4$ for AP-1 and $\chi2=5.5$, $p=<0.06$ for lola-PT.

| Genotype | Average Age | Median Age | n | p value vs outcrossed GAL4 | p value vs outcrossed transgene |
|---|---|---|---|---|---|
| MHC-GAL4::w1118 | 3.8 days | 3 days | 75 | n/a | n/a |
| w1118::AP-1 | 5.7 days | 5 days | 145 | 4.80E-05 | n/a |
| MHC-GAL4::AP-1 | 7.5 days | 8.5 days | 120 | 8.10E-13 | 2.50E-09 |
| w1118::lola PT | 5.2 days | 5 days | 106 | 0.01258 | n/a |
| MHC-GAL4::lola PT | 5.7 days | 6 days | 123 | 0.00085 | 0.06162 |

**Table S8.** Survival of AP-1 and lola-PT over-expression with 20mM paraquat exposure. $\chi 2=70$, $p=6E-16$ for AP-1 and $\chi 2=15.8$, $p=4E-04$ for lola-PT.

| Genotype | Average Age | Median Age | n | p value vs outcrossed GAL4 | p value vs outcrossed transgene |
|---|---|---|---|---|---|
| MHC-GAL4::w1118 | 11.5 days | 12 days | 66 | n/a | n/a |
| w1118::AP-1 | 14.3 days | 14 days | 58 | 0.03915 | n/a |
| MHC-GAL4::AP-1 | 16.9 days | 17.5 days | 90 | 1.80E-09 | 0.00043 |
| w1118::lola PT | 10.7 days | 11 days | 68 | 0.00067 | n/a |
| MHC-GAL4::lola PT | 14.6 days | 13 days | 98 | 0.00116 | 5.30E-07 |

**Table S9.** Survival of AP-1 and lola-PT over-expression with 10mM paraquat exposure. $\chi2$=35.1, p=2E-08 for AP-1 and $\chi2$=33.5, p=5E-08 for lola-PT.

| Species | Average Age | Median Age | n | Confidence Group |
|---|---|---|---|---|
| *D. melanogaster* | 7.5 days | 8 days | 210 | a |
| *D. mauritiana* | 9.4 days | 8 days | 207 | e |
| *D. ananassae* | 5.7 days | 6 days | 212 | b |
| *D. simulans* | 9.8 days | 10 days | 211 | e, f |
| *D. pseudoobscura* | 15.3 days | 16 days | 200 | c |
| *D. virilis* | 12.8 days | 11 days | 245 | d |
| *D. yakuba* | 9.2 days | 9 days | 209 | e, g |

**Table S10.** Survival of *Drosophila* species with 10mM paraquat exposure. $\chi 2=547$, $p=<2e-16$.

| Binding Site | Interaction p-value |
|---|---|
| lola-PO | p=0.14149 |
| motif 4 | p=0.10614 |
| motif 5 | p=0.52630 |

**Table S11.** Interaction plots between levels of p38K and the presence of either the lola-PO, motif 4, or motif 5 binding sites.
